# Engineering immunotoxin-equipped effector cells and evaluation in primary human immune cells

**DOI:** 10.64898/2025.12.09.691975

**Authors:** Alexander H. Pearlman, Brian J. Mog, Michael S. Hwang, Jordina Rincon-Torroella, Sarah R. DiNapoli, Suman Paul, Jacqueline Douglass, Emily Han-Chung Hsiue, Stephanie A. Glavaris, Drew M. Pardoll, Nickolas Papadopoulos, Kenneth W. Kinzler, Chetan Bettegowda, Shibin Zhou, Bert Vogelstein, Maximilian F. Konig

**Affiliations:** Ludwig Center and Lustgarten Laboratory, Sidney Kimmel Comprehensive Cancer Center, Johns Hopkins University School of Medicine, Baltimore, MD 21287; Howard Hughes Medical Institute, Chevy Chase, MD 20815; Department of Neurosurgery, Johns Hopkins University School of Medicine, Baltimore, MD 21287; Division of Hematologic Malignancies and Bone Marrow Transplantation, Department of Oncology, Johns Hopkins University School of Medicine, Baltimore, MD 21287; Bloomberg∼Kimmel Institute for Cancer Immunotherapy, Johns Hopkins University School of Medicine, Baltimore, MD 21287; Department of Oncology, Johns Hopkins University School of Medicine, Baltimore, MD 21287; Department of Pathology, Johns Hopkins University School of Medicine, Baltimore, MD 21287; Institute for NanoBioTechnology, Johns Hopkins University, Baltimore, MD 21287; Division of Rheumatology, Department of Medicine, Johns Hopkins University School of Medicine, Baltimore, MD 21287

## Abstract

Lethal toxins could become potent therapies against cancer, but their clinical utility is limited by adverse events upon systemic administration. These could be reduced if the toxins were delivered by effector cells that specifically infiltrate cancers, thereby releasing toxins locally into the tumor microenvironment. One of the challenges underlying this strategy is that cells delivering toxins would have to be resistant to them. We address this obstacle by showing that effectors derived from transformed human cell lines genetically engineered for resistance to bacterial adenosine diphosphate ribosylating toxins (ADPRTs), including *Pseudomonas aeruginosa* exotoxin A (PE), can produce targeted immunotoxins that specifically kill cancer cells expressing cognate tumor-associated antigens. Resistance to immunotoxins was achieved by knockout of genes in the diphthamide biosynthesis pathway (*DPH1-4*) required for the posttranslational modification of eukaryotic elongation factor 2 (EEF2) that is the target of ADPRTs, or by mutation of *EEF2* itself. We show that engineering resistance to ADPRTs, one of the most potent toxins acting on human cells, is essential to achieve robust function of armored effector cell lines. This work establishes a first step on the path to equip effector cells with the ability to deliver powerful toxins to cancer cells and introduces a platform to investigate extension to primary autologous or allogeneic therapeutic cell types.

## Introduction

Immunotoxins, recombinant proteins comprising a targeting domain fused with a bacterial or plant toxin, can be powerful cancer therapeutics. Three immunotoxins have received regulatory approval for clinical use in the United States, showing efficacy in cutaneous T-cell lymphoma (interleukin-2-immunotoxin), blastic plasmacytoid dendritic cell neoplasm (CD123-targeted immunotoxin), and hairy cell leukemia (CD22-targeted immunotoxin)^1–7^. Despite these successes, the widespread use of immunotoxins has been constrained by their toxicities upon systemic administration and by the development of anti-drug antibodies^8–11^. Engineered cellular therapies, such as chimeric antigen receptor (CAR)-T cells, have similarly achieved remarkable success against leukemias and lymphomas, but have so far been less efficacious against solid tumors. The heterogeneity of antigen expression in tumors is believed to be one of the main barriers limiting CAR-T cell effectiveness^12–15^. Recent clinical evidence has indicated that CAR-T cells secreting bispecific antibodies to another antigen might be able to overcome this obstacle^16^. However, a second barrier, intrinsic tumor cell resistance to perforin/granzyme-, tumor necrosis factor-α (TNFα)-, or interferon-γ (IFNγ)-mediated cytotoxicity, would not be overcome, even in theory, by local secretion of bispecific antibodies^17–21^.

Engineering T cells to express a synthetic immune receptor (e.g., CAR) and secrete a suitable immunotoxin could overcome the limitations of these two therapeutic modalities. Local secretion of immunotoxin by CAR-T cells infiltrating the tumor could reduce or eliminate systemic toxicities of the immunotoxin. This local secretion could also provide a bystander effect, killing cancer cells or stromal cells of the tumor microenvironment that do not express the antigen to which the CAR-T cells are directed. Importantly, immunotoxins kill tumor cells via an orthogonal mechanism to immune effector cell cytotoxicity pathways, thus overcoming acquired resistance to perforin/granzyme-mediated cytotoxicity, the second barrier noted above. Finally, immunotoxin activity may induce an inflammatory tumor microenvironment that would enhance CAR-T cell activity^22–26^.

Most immunotoxins, including those approved for clinical use, are derived from either exotoxin A of *Pseudomonas aeruginosa* (PE) or diphtheria toxin of *Corynebacterium diphtheriae* (DT). These bacterial toxins catalyze the adenosine diphosphate (ADP)-ribosylation of eukaryotic elongation factor 2 (EEF2) at its diphthamide-modified histidine residue (H715), a post-translational modification thought to be unique to EEF2^27^. Toxin-induced ADP-ribosylation of EEF2 then leads to irreversible arrest of protein synthesis and secondary cell death^28,29^. Any mammalian cell, including CAR-T cells secreting immunotoxins, would be susceptible to ADP-ribosylating toxins (ADPRTs) if their catalytic domains entered the cytosol. Therefore, the effective production of immunotoxins by CAR-T cells requires engineered resistance to ADPRTs. Previous studies have reported producing immunotoxins in cells with resistance derived by spontaneous mutagenesis or transiently in non-resistant immune cells^30–40^. The implementation of a defined, engineerable resistance mechanism that allows engineered cells to produce immunotoxins is an essential first step for advancing immunotoxin-armored cellular therapies for the treatment of cancer. Disruption of the diphthamide biosynthesis pathway or introduction of an *EEF2* p.G717R mutation yields cells that are genetically resistant to toxin activity^29,41–45^. Here, we describe the creation of human effector cells from transformed cell lines, including Jurkat T cells, that can secrete immunotoxins with specific activity against cancer cells while maintaining engineered resistance (Fig. 1).

**Figure 1:**
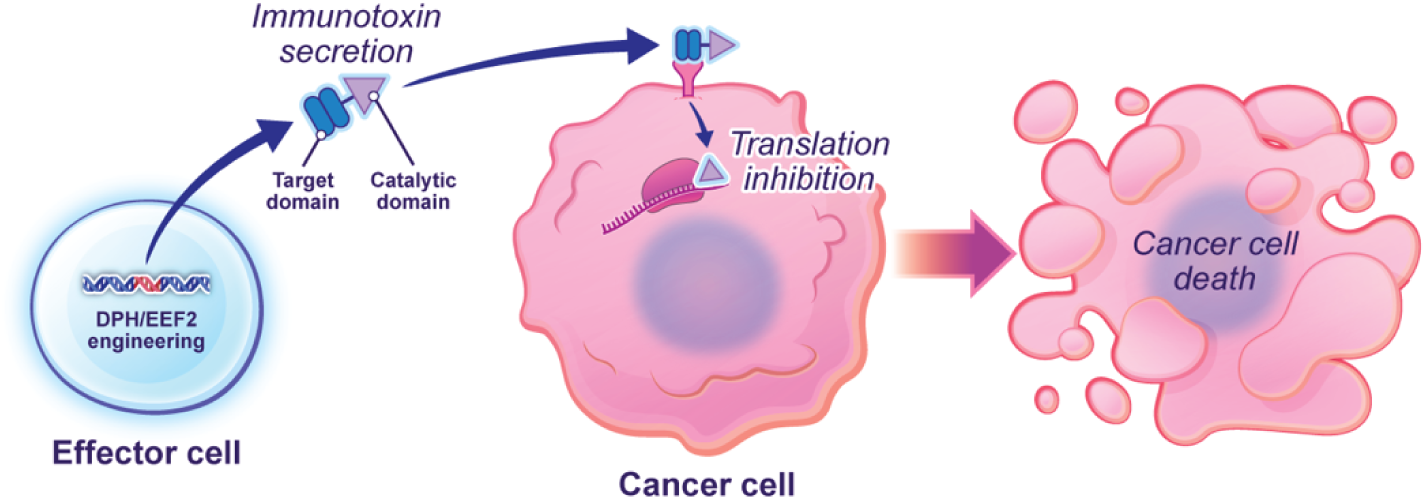
Engineered effector cells secreting immunotoxins as a cancer therapeutic. Effector cells are engineered via knockout of diphthamide biosynthesis genes or EEF2 mutation to be resistant to ADPRT immunotoxins. The effector cells locally secrete immunotoxins in tumors, leading to inhibition of translation and subsequently cancer cell death.

## Results

### Engineering cellular resistance to immunotoxins

To efficiently express and secrete immunotoxins in human effector cells, we first engineered cellular resistance to ADPRTs. 293FT cells were treated with recombinant, full-length DT or PE, and cell survival was measured at various times thereafter. Mammalian 293FT cells were sensitive to both DT and PE, with intoxication leading to cell death (Fig. 2A-B). Next, we used CRISPR/Cas9 to disrupt known genes of the diphthamide biosynthesis pathway in 293FT cells using guide RNAs (gRNAs) to selectively target *DPH1*, *DPH2*, *DPH3*, *DPH4*, *DPH5*, *DPH6*, or *DPH7* (Supplementary Table S1 and Supplementary Fig. S1). The inactivation of any of these genes was expected to prevent biosynthesis of the diphthamide modification of EEF2 at amino acid H715, the shared catalytic target of ADPRTs. In an orthogonal approach, we used CRISPR/Cas9 homology-directed repair to introduce a p.G717R mutation in the endogenous *EEF2* gene locus in 293FT cells (Supplementary Table S1 and Supplementary Fig. S1). This mutation at amino acid 717 is known to abrogate diphthamide modification of the nearby (2 amino acid N-terminal) H715^46^. Next, we treated the genetically modified 293FT cells with exogenous DT and measured cell growth over time. Disruption of the *DPH1*, *DPH2*, *DPH3*, and *DPH4* genes, as well as introduction of mutant EEF2 p.G717R, conferred resistance to DT and PE (Fig. 2C-D and Supplementary Fig. S2). Although *DPH3* disruption in 293FT cells resulted in resistant cells, edited cells demonstrated reduced fitness over time when grown in the presence of DT. 293FT cells with *DPH5* disruption displayed a reduced rate of growth in the presence of DT relative to other mutants. *DPH6* and *DPH7* disrupted cells did not remain viable in the presence of DT, indicating either incomplete toxin resistance or reduced fitness. Knockout of *DPH1* in a transformed human T cell line (Jurkat) (Fig. 2E) or in primary human T cells (Fig. 2F) similarly rendered cells resistant to exposure to exogenous DT, which invariably resulted in arrest of cell division and death in unedited T cells. This established that knockout of a subset of *DPH* genes as well as introduction of an *EEF2* p.G717R mutation can confer ADPRT resistance to effector cells expressing diphtherial immunotoxin.

**Figure 2:**
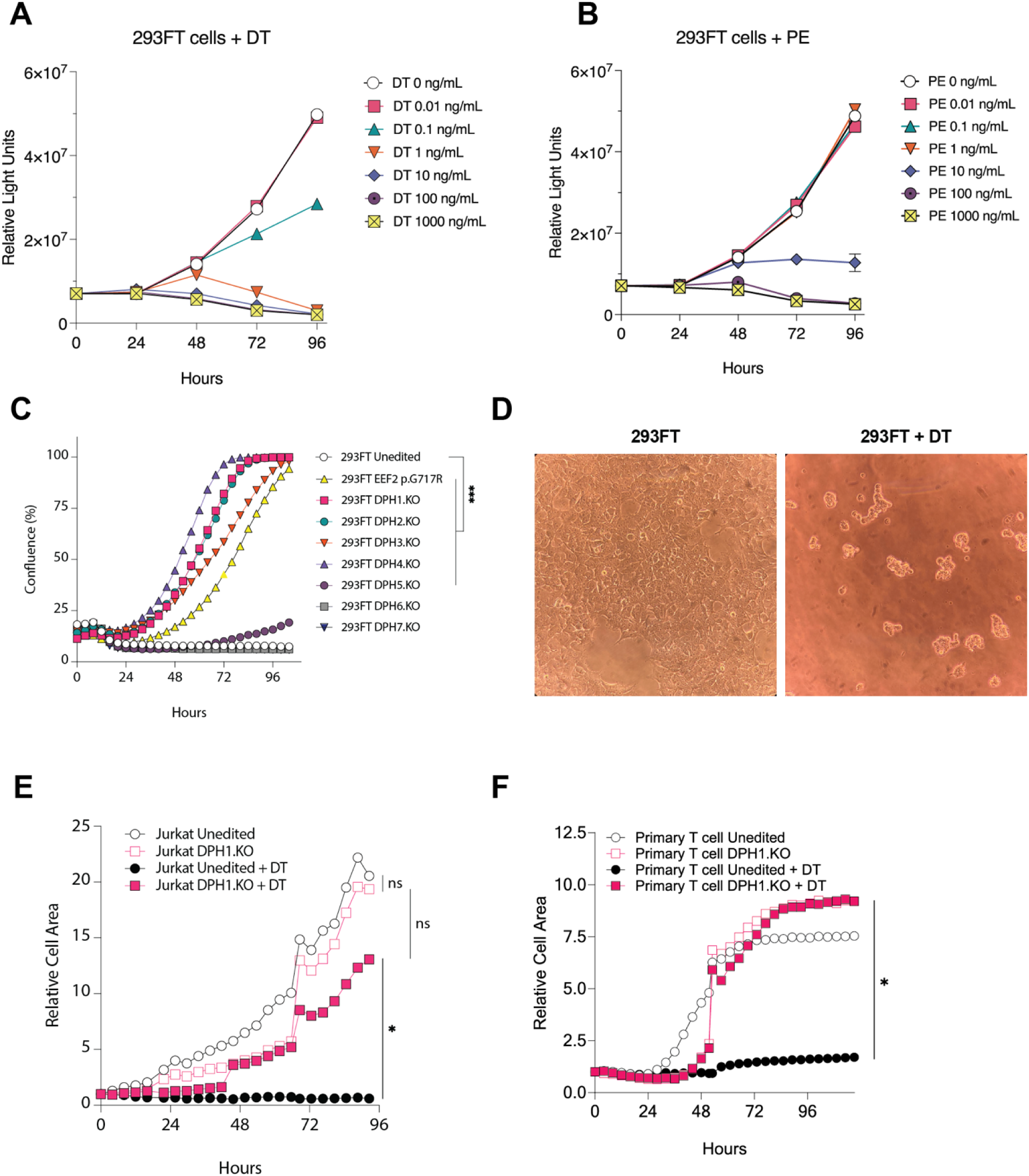
Engineering ADP-ribosylating toxin resistance in mammalian cells through disruption of DPH genes and EEF2 mutagenesis. (A, B) Growth of 293FT cells cultured in the presence of varying concentrations of full-length, bacterial DT (A) or PE (B) was quantified at various time points using the CellTiterGlo luciferase activity assay. (C) Unedited 293FT cells, 293FT cells edited with CRISPR/Cas9 to disrupt *DPH1-7* (DPH1.KO-DPH7.KO), or 293FT cells edited to introduce mutant EEF2 p.G717R were cultured in the presence of 1000 ng/mL native DT. Cell confluence over time was quantified using time-lapse microscopy. ***: p<0.001 by one-way ANOVA, Dunnet’s test. (D) Morphology of 293FT non-engineered cells by light microscopy at 10X magnification in the absence or presence of DT. DT arrests cell proliferation and induces characteristic morphological changes and secondary cell death. (E, F) Jurkat cells (E) and primary human T cells (F), either unedited or edited to knockout *DPH1* (DPH1.KO), were cultured in the absence or presence of 2000 ng/mL DT. Relative cell area over time was quantified using live-cell microscopy. ns: not significant, *: p<0.05 by Kruskal-Wallis test with Dunn’s test. DT: *Corynebacterium diphtheria* toxin. PE: *Pseudomonas aeruginosa* exotoxin A.

### T cells with engineered resistance to immunotoxins maintain effector functions

Previous reports have shown that disruption of diphthamide biosynthesis genes can impair cell growth in yeast^47^. Impaired replication could be detrimental to human immune effector cells, which generally rely on in-vivo expansion for efficacy^48^. To evaluate this possibility, we investigated the functional properties of modified effector cells in vitro. We used CRISPR/Cas9 to disrupt *DPH1*, *DPH2*, *DPH3*, or *DPH4* or to introduce an *EEF2* p.G717R mutation in primary human T cells. Edited T cells were then expanded in the presence of exogenous, full-length DT for 9 days to select for resistant T cells. Engineered T cells proliferated under DT exposure whereas unedited cells did not (Fig. 2E, Supplementary Fig. S3). We then co-cultured edited T cells (after removal of DT from media) and unedited T cells (not exposed to DT) with CD19+ NALM6 B cells in the presence or absence of a CD19xCD3 bispecific T cell engager, to activate T cells and induce T cell-mediated killing of NALM6 target cells^49^. Disruption of *DPH* genes or *EEF2* mutation did not prevent T cell proliferation (Fig. 3A and Supplementary Fig. S4), cytokine production (Fig. 3B), or T cell-mediated cytotoxicity (Fig. 3C), and allowed effective killing of NALM6 cells in vitro.

**Figure 3:**
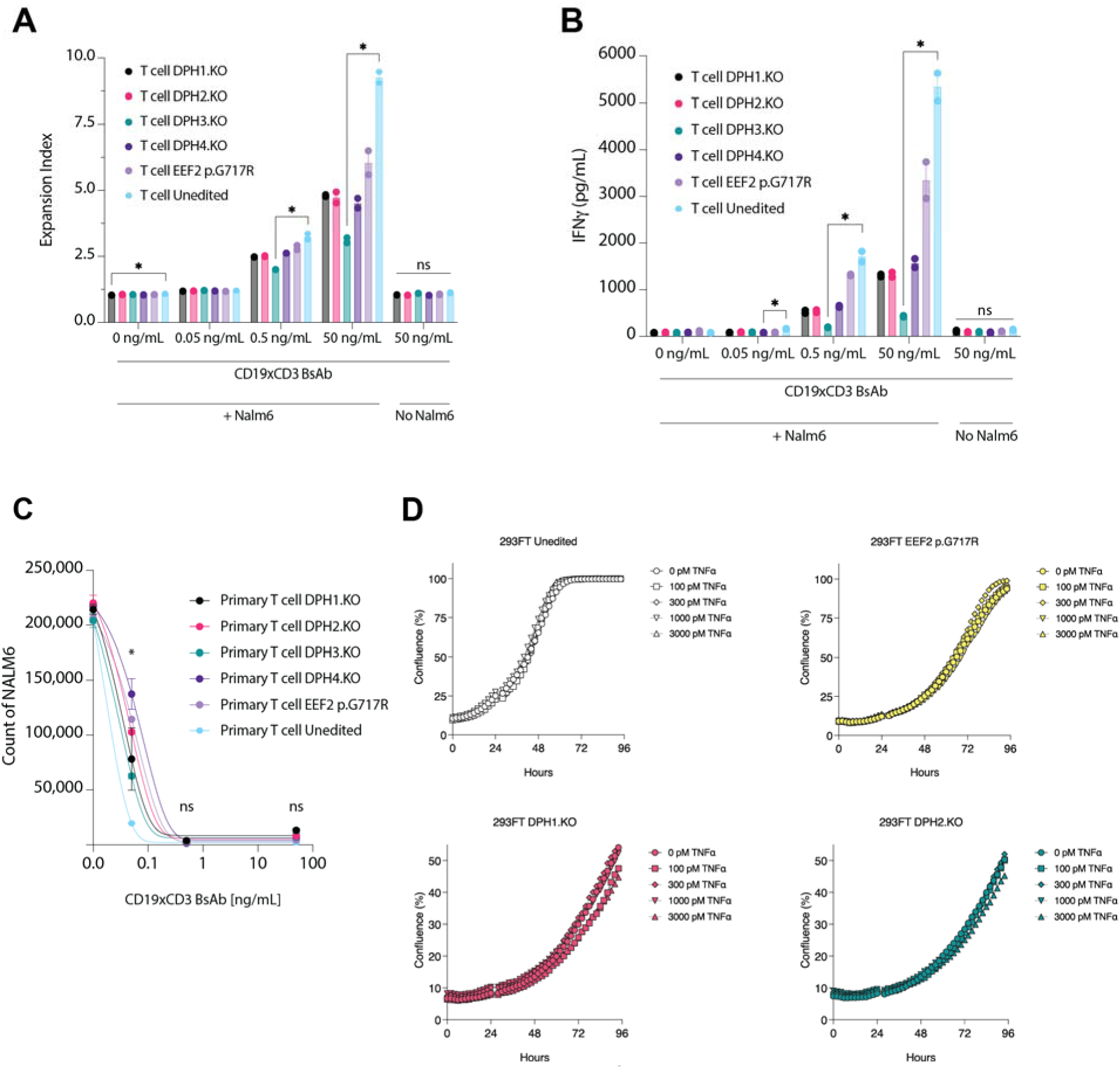
Primary human T cells engineered for APPRT resistance maintain effector functions for cancer immunotherapy. (A, B, C) CRISPR/Cas9 was used to knockout *DPH1*, *DPH2*, *DPH3*, or *DPH4* or to introduce an *EEF2* p.G717R mutation in primary human T cells to confer ADPRT resistance. After selection in DT, edited T cells were co-cultured with CD19+ NALM6 target cells and a CD19xCD3 bispecific T cell-engaging antibody (BsAb) at 0, 0.05, 0.5, or 50 ng/mL. T cell expansion index (A), IFNγ secretion in supernatant (B), and NALM6 cell counts (C) were quantified by flow cytometry on day 7. ns: not significant; *: p<0.05 by Kruskal-Wallis test with Dunn’s test. (D) 293FT cells edited with CRISPR/Cas9 for *DPH1* or *DPH2* knockout or *EEF2* p.G717R mutation were selected in DT and then cultured in the presence of exogenous TNFα. Cell confluence over time was quantified by live cell microscopy. BsAb: bispecific antibody. DT: *Corynebacterium diphtheria* toxin. IFNγ: interferon-γ. TNFα: tumor necrosis factor-α.

Additionally, prior studies have reported that diphthamide biosynthesis pathway disruption can predispose cells to TNFα-mediated apoptosis^41^. If this effect were pronounced in human effector cells, this would be problematic for cellular therapies where TNF action is a key mediator of tumor cell cytotoxicity^17^. To assess sensitivity to TNFα, we disrupted *DPH1* or *DPH2* or introduced mutant *EEF2* p.G717R in 293FT cells. After selection for ADPRT resistant edited cells with exogenous DT, we cultured resistant 293FT with human TNFα and measured cell growth over time (Fig. 3D). We found that TNFα exposure did not meaningfully decrease growth rates in cells with *DPH* gene disruption or *EEF2* mutation compared to their original counterparts. In addition, engineered resistance did not lead to increased 293FT cell death compared to non-engineered cells under exposure to high TNFα concentrations (Supplementary Fig. S5). These results collectively suggested that using *DPH* gene knockout or *EEF2* mutation as a resistance mechanism would not greatly impair effector cell activity and growth.

### Immunotoxins secreted by engineered effector cells induce target cell death

With a resistance mechanism established, we proceeded to evaluate candidate immunotoxin constructs to determine which ones could be effectively expressed by mammalian cells (Supplementary Table S2 and Supplementary Fig. S6). For this screen, candidate immunotoxins were targeted against human epidermal growth factor receptor (EGFR) using either a single chain variable fragment (scFv) derived from the anti-EGFR antibody cetuximab or truncated epidermal growth factor (tEGF) as a ligand for EGFR. These targeting domains were fused to several toxin variants derived from the catalytic and translocation domains of DT or PE^50–54^. After transfecting immunotoxin constructs into various resistance-modified 293FT cells, we measured immunotoxin protein expression by staining for DYKDDDDK (FLAG)-tag using intracellular flow cytometry. GFP constructs were used as transfection controls (Supplementary Fig. S7). A bispecific antibody with a FLAG-tag was used as a control for transfection and staining^55^. We observed that all immunotoxin constructs could be expressed in ADPRT-resistant 293FT cells at levels similar to that of a cetuximab scFv control (Fig. 4A). Viable immunotoxin expressing cells were depleted when active immunotoxin constructs were transfected into original, non-resistant 293FT cells, indicating that all fusion toxins were indeed actively cytotoxic (Fig. 4A). At the same time, transfection of inactive mutant immunotoxin constructs (DTΔ180-cetux, cetux-PE38Δ578) into non-resistant 293FT cells yielded viable cells producing the constructs (Fig. 4A).

**Figure 4:**
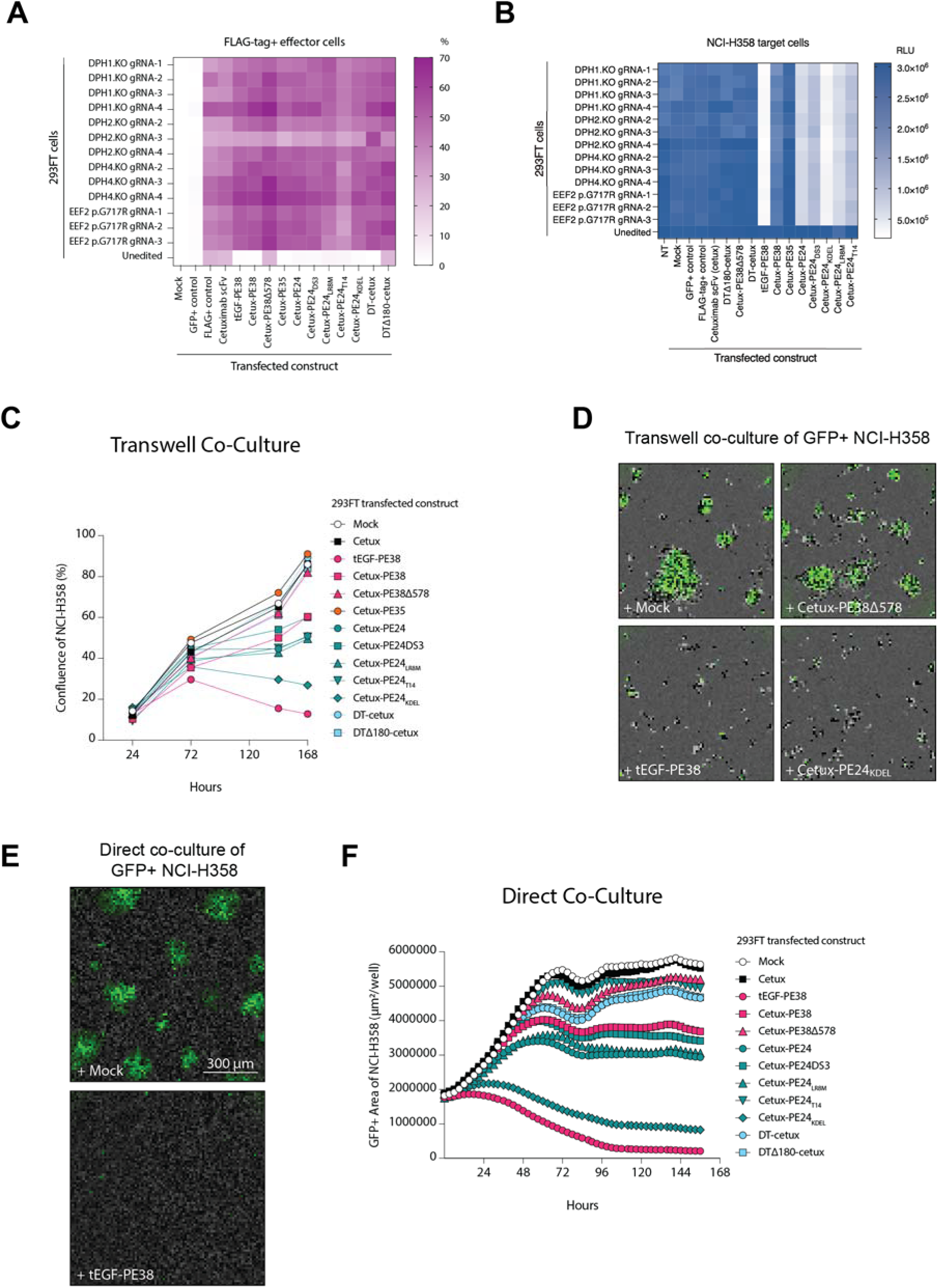
Immunotoxins expressed by engineered 293FT cells are biologically active and induce cancer cell death. (A) 293FT cells engineered for resistance with knockout of *DPH* genes (DPH.KO) or EEF2 p.G717R mutation were transfected for expression of immunotoxin or control constructs. The percentage of live cells expressing the constructs was quantified by intracellular flow cytometry staining of FLAG-tag expressed as part of the constructs. DTΔ180 and PE38Δ578 are catalytically inactive forms of DT and PE fusion proteins, respectively. Mock: 293FT cells subjected to mock transfection protocol with no DNA. (B) Conditioned media supernatant from transfected 293FT cells was added to NCI-H358 cells. Cancer cell viability of NCI-H358 was measured by SteadyGlo luciferase activity after 4 days of supernatant exposure. NT: non-transfected. (C) 293FT cells engineered for ADPRT resistance with *DPH2* knockout were transfected with immunotoxin constructs and then co-cultured across a transwell insert from NCI-H358 target cells. Confluence of NCI-H358 cells was quantified over time using live cell microscopy. (D) Example images of GFP+ NCI-H358 cells in transwell co-culture with resistance-engineered 293FT cells transfected in the absence of DNA construct (Mock), or with a DNA construct encoding inactive mutant cetuximab-PE38 (+Cetux-PE38Δ578), encoding active tEGF-PE38 (+tEGF-PE38), or encoding active cetuximab-PE24_KDEL_ (+Cetux-PE24_KDEL_). Images were acquired on day 5 of co-culture at 10X magnification. (E) Example images of GFP+ NCI-H358 cells in direct co-culture with resistance-engineered 293FT cells transfected in the absence of DNA construct (Mock) or with DNA encoding tEGF-PE38 (+tEGF-PE38). Representative images acquired at 10X magnification on day 5 of co-culture are shown. 293FT cells are GFP-negative and observed in the background. (F) 293FT *DPH1* knockout cells were transfected with various immunotoxin constructs and then directly co-cultured with NCI-H358 cells. GFP-labeled NCI-H358 cells were quantified longitudinally using live cell microscopy. NT: non-transfected.

To evaluate the secretion and function of the immunotoxins, we transfected the same constructs into resistance-modified 293FT cells and tested the activity of the EGFR-targeted immunotoxins on NCI-H358 target cells. NCI-H358 expresses EGFR and is derived from a patient with lung cancer. We engineered them to express eGFP and luciferase to more easily follow their growth. Culture media from resistance-engineered 293FT cells transfected to express immunotoxins were collected and cleared of cells by centrifugation and the supernatant was then applied to the NCI-H358 target cells. Six resistance-engineered cell lines transfected with immunotoxin constructs yielded supernatants that induced substantial cell death in the NCI-H358 target cells as measured by target cell luciferase activity or confluence (Fig. 4B and Supplementary Fig. S8). Target cells treated with supernatants containing active immunotoxins exhibited the morphologies characteristic of ADPRT intoxication and ADPRT-induced cell death (Supplementary Fig. S9)^56,57^. Negative controls for this experiment included ADPRT-resistant cells transfected with several constructs that did not include toxin modules, as well as supernatants from 293FT cells that had not been made resistant to ADPRT (Fig. 4B and Supplementary Fig. S8). Three (DT-cetuximab, cetuximab-PE38, and cetuximab-PE35) of the nine toxin-containing constructs did not lead to supernatants that killed target cells, suggesting that those immunotoxins were not secreted, were secreted in an inactive form, or were not internalized by target cells.

To ensure that the immunotoxins were secreted in soluble form, we co-cultured transfected 293FT cells across the barrier insert of transwell plates containing NCI-H358 cells in the lower chamber (Fig. 4C-D and Supplementary Fig. S10). The relative potency of the constructs observed in Fig. 4B was largely preserved in the transwell experiments. We also directly co-cultured 293FT cells transfected with various immunotoxins constructs with NCI-H358 cells and followed the time course of target cell death by time lapse microscopy (Fig. 4E-F and Supplementary Video S1-S2). Cytotoxicity was observed using effectors with both DPH knockout (Fig. 4C, F and Supplementary Fig. S11A)

To evaluate other immunotoxins in this system, we generated constructs incorporating the EGFR-targeting scFvs 806 or D2C7ds instead of tEGF or the cetuximab domain recognizing the EGFR antigen (Supplementary Table S2, Supplementary Fig. S11B-C, Supplementary Fig. S12)^58,59^. We also tested a third bacterial ADPRT toxin, derived from the catalytic domain of *Vibrio cholerae* exotoxin (CET) (Supplementary Table S2, Supplementary Fig. S11B-C, Supplementary Fig. S12)^60^. Overall, we observed that the tEGF-PE38 immunotoxin, of all tested immunotoxin designs, had the greatest cytotoxicity in these assays (Supplementary Fig. S11D).

### Resistance to immunotoxins in model T cells is required for target cell killing

We next used CRISPR/Cas9 to disrupt *DPH1* in Jurkat T cells, serving as a T cell model, and thereby confer toxin resistance. We then transfected the engineered Jurkat cells with a DNA construct to express the tEGF-PE38 immunotoxin. To test the specificity of the secreted immunotoxin, we co-cultured the immunotoxin-equipped Jurkat cells with a variety of EGFR+ and EGFR- target cells and quantified the growth of target cells using time-lapse microscopy. Jurkat cells secreting the tEGF-PE38 immunotoxin did not kill ADPRT-resistant target cells nor cells that do not express EGFR, as expected (Fig. 5A). However, they did kill several patient-derived target cell lines that express EGFR (Fig. 5A). Finally, we compared the cytotoxic ability of the original (non-ADPRT-resistant) Jurkat cells with their ADPRT-resistant counterparts after transfection with the tEGF-PE38 immunotoxin construct. Unlike the ADPRT-resistant Jurkat T cells, the original Jurkat cells transfected with immunotoxin constructs exhibited no toxicity against target cells expressing the cognate EGFR antigen (Fig. 5B-C). Similarly, introducing

**Figure 5:**
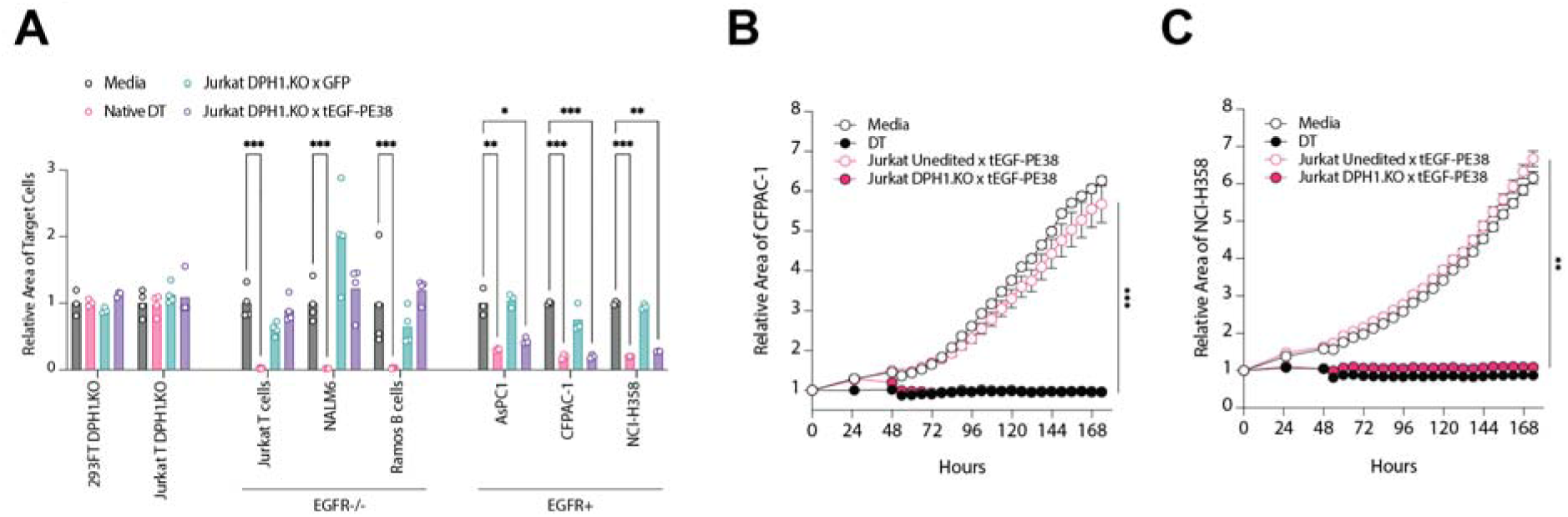
Resistance-engineered Jurkat T cells equipped with immunotoxin induce selective killing of cancer cells. Target cells were co-cultured across a transwell barrier with media alone, exogenous DT, Jurkat DPH1.KO effector T cells transfected to express GFP, or Jurkat DPH1.KO effector T cells transfected to express tEGF-PE38. Target cell area across time was quantified. *: p<0.05; **: p<0.01; ***: p<0.001 by two-way ANOVA with Dunnett’s test. (B, C) Jurkat cells without ADPRT resistance or Jurkat DPH1.KO cells with resistance were transfected to express tEGF-PE38 and co-cultured across a transwell barrier with CFPAC-1 (B) or NCI-H358 (C) target cells. Target cell area across time was quantified. **: p<0.05; ***: p<0.001 by one-way ANOVA with Tukey’s test. DT: *Corynebacterium diphtheria* toxin.

### Evaluation of application in primary human immune cells

With the basis established in transformed human cells, we then evaluated extension of the strategy to primary human immune cells, a potentially more therapeutically relevant platform. We engineered primary human T cells for ADPRT resistance using CRISPR and then transduced them with lentiviral vectors of control or immunotoxin constructs. These immunotoxin constructs also contained as an expression marker a co-translated CAR (V2.CAR) with a targeting domain derived from a single-chain variable fragment (scFv) that recognizes a *KRAS* p.G12V neoantigen peptide presented by HLA-A*03^55^. After transduction, we co-cultured the T cells with NCI-H358 target cells. Unlike Jurkat T cells tested previously, primary T cells exhibited no cytotoxicity against NCI-H358 cells (Supplementary Fig. S13A, B). When constructs were introduced using a second round of CRISPR knockin, <1% of T cells expressed the immunotoxin construct (Supplementary Fig. S13C). To test alternative expression methods and obviate the need for transcription, we transfected resistance engineered primary T cells with mRNA through electroporation or with mRNA lipid nanoparticles. These T cells again did not exhibit cytotoxicity when co-cultured with NCI-H358 target cells (Supplementary Fig. S13D, E). To attempt to more finely control potential expression from immunotoxin constructs, we used CRISPR knockin to introduce into T cells an inverted immunotoxin encoding sequence that could not be transcribed. We then treated T cells with Cre recombinase to recombine the knocked in constructs into an orientation compatible with transcription. These constructs showed inconsistent and low expression of a linked CAR marker (Supplementary Fig. S13F).

Because we did not observe expression in primary T cells, we evaluated the RNA constructs in transformed cell lines as comparison. The constructs exhibited cytotoxicity when directly transfected into non-resistant NCI-H358 cells but not when transfected into resistance-engineered 293FT cells (Supplementary Fig. S14A). Strong expression of constructs was further apparent with flow cytometry measurement of protein L staining of the CAR marker scFv or with intracellular staining of an attached FLAG-tag (Supplementary Fig. S14B, C). Comparing across cell types and expression modalities, immunotoxin component expression could be detected only with transfection of 293FT cells and not with lentiviral transduction of 293FT cells nor transduction of primary T cells despite evident expression of the linked CAR marker (Supplementary Fig. S14D, E). However, there did appear to be at least some expression in primary T cells of immunotoxin components below the limit of detection of our assays as non-resistant primary T cells transfected with CAR-immunotoxin constructs showed no CAR component expression whereas resistance-engineered primary T cells did (Supplementary Fig. S14F, G).

To rule out any mechanisms specifically related to PE that could be limiting immunotoxin expression in primary T cells, we then tested immunotoxin designs based on DT rather than PE catalytic domains. The DT-based constructs showed expected cytotoxicity when transfected into non-resistant NCI-H358 cells but not when transfected into resistant 293FT *DPH* knockout cells (Supplementary Fig. S15A). Primary T cells transfected to express these constructs again showed no cytotoxicity in co-culture with NCI-H358 targets (Supplementary Fig. S15B).

We finally evaluated whether cell-intrinsic factors could be constraining primary T cell immunotoxin expression. To test if furin activity could be influencing immunotoxin expression, we knocked out furin in T cells together with *DPH1*. Expression as measured with cytotoxicity in co-culture did not appear to be changed (Supplementary Fig. S16A). Primary human B cells engineered for ADPRT resistance and transfected with immunotoxin constructs showed no cytotoxicity in co-culture with NCI-H358 target cells(Supplementary Fig. S16B). Similar results as in primary T cells were also observed with NALM6 (B cell derived) and NK-92 (NK cell derived) transformed cell lines and with B-lymphoblastoid cell lines (B-LCL) generated from primary B cells (Supplementary Fig. S16C, D). In primary NK cells, expression of linked FLAG-tag as measured with flow cytometry was similarly limited (Supplementary Fig. S16E).

## Discussion

The experiments reported herein lay the foundation for therapeutic approaches in which effector cells can be equipped with the ability to secrete molecules that are highly toxic to cancer cells. A key element of this study is the demonstration that resistance to the toxic agent must be engineered in effector cells to be used as therapeutic agents. With highly potent protein toxins, such as the immunotoxins used here, therapeutic effector cells will themselves be rapidly killed or disarmed by the toxin payload if this resistance is not engineered into them. Immunotoxins, or other proteins that directly kill cancer cells, have advantages over other proteins that can be secreted by the effector cells. For example, bispecific T cell-engaging antibodies secreted by CAR-T cells have been shown to have bystander effects and augment the cytotoxic activity of the engineered immune effector cells^14,16^. This augmentation, however, is dependent on the presence of abundant T cells within the tumor, avoidance of local immunosuppression or T cell exhaustion, and preserved susceptibility of the tumor to perforin/granzyme-mediated cytotoxicity. In contrast, immunotoxins derived from bacterial ADPRTs do not require T cells to kill their target cells, induce cell death by an orthogonal perforin/granzyme-independent mechanism (arrest of protein synthesis), curtail repair and escape mechanisms of tumor cells by irreversibly arresting protein biosynthesis, and are the most efficient cytotoxic agents that have ever been discovered. It is estimated that even a single toxin molecule reaching the cytosol of a target cell may be sufficient to kill it^61^.

In our studies, we successfully expressed immunotoxins with a variety of combinations of targeting domains and catalytic domains. Targeting domains included either antibody fragments (e.g., scFvs of cetuximab, 806, D2C7ds) or ligands (tEGF)^58,59^. Catalytic domains used in our immunotoxins were derived from DT, PE, or the less commonly studied CET^50–53,60^. In addition, we used different fragments of PE toxin (e.g., PE38, PE24), toxin domains modified to protect the furin cleavage site (PE24_DS3_), to increase endoplasmic reticulum trafficking with the KDEL receptor (PE24_KDEL_), or to reduce immunogenicity (PE24_LR8M_, PE24_T14_). These variations reflect the diversity of targeting and catalytic domains in clinically approved immunotoxins and those in preclinical development^1–3^. Despite strong expression, not all immunotoxin varieties demonstrated strong cytotoxic activity in our models. We observed strongest activity from our cetux-PE24_KDEL_, tEGF-PE38, and tEGF-CET40 designs, while DT-based immunotoxin showed cytotoxicity in unmodified effector cells but not on target cells. These findings suggest that determining optimal combinations of toxin domains and binding moieties for secretion by effector cells requires empirical testing.

Our study provides a platform to systematically evaluate factors controlling the performance of immunotoxins delivered by human effector cells, which may be different from those influencing traditional immunotoxins that are produced in bacteria and delivered exogenously. Traditional considerations driving the optimization of immunotoxins for systemic administration, including immunogenicity and off-target toxicity, may not be primary constraints for therapies that are locally and conditionally secreted in the tumor environment. Instead, design optimization must focus on abolishing innate cellular responses to the immunotoxin by secreting cells and increasing the efficiency of immunotoxin expression and secretion.

The approach described here is general, as we have shown it can be applied to several cell types as delivery vehicles as well as to several immunotoxins. Many different cell types, ranging from dendritic cells to tumor cells themselves, have been proposed to be potentially useful as therapeutic agents^62,63^. But, we emphasize that this study represents only the first step towards achieving a clinically useful cell-based therapeutic agent that leverages immunotoxins for bystander killing. Optimal future approaches would incorporate transcriptional signaling into the immunotoxin secretion, so that inducible secretion would only be achieved once the effector cell mediating the delivery is within the tumor. This transcriptional activation could be downstream of CAR signaling or initiated by other types of synthetic receptors^64^. The cells would also ideally be long-lasting, as expansion and persistence of effector cells has been shown to be critical for tumor control and elimination in some settings^65–67^.

To date, we have been unable to stably express immunotoxins in any of the cells described herein, either using lentivirus transduction or CRISPR-Cas-mediated gene knock-in. Although we have been able to achieve potent transient expression in cultured transformed cells, we have not been able to achieve transient expression in ADPRT resistance-engineered primary T cells, B cells, or NK cells. The discrepancy between transformed cells and primary cells suggests a critical biological difference in cellular responses to immunotoxins whose mechanisms have not been discerned to date. Because all the tested immunotoxin designs contain a furin cleavage site, the cleavage of which is important for toxin trafficking, one important difference may be the level of furin expression in the different cell types. T cells have high levels of furin expression, upregulated upon activation, whereas 293T cells have low levels of furin expression^68,69^. Differential levels of furin may therefore lead to differential furin cleavage, affecting toxin trafficking and expression. Similarly, differences in endoplasmic reticulum associated degradation could be limiting toxin expression^70^. Another explanation could involve toxin activation of innate immunity signaling pathways through the inflammasome, leading to effector cell death^71^. Our current work is devoted to understanding and overcoming the barriers that have limited stable expression in primary cells. This will allow us to expand from the critical steps reported here to achieve persistently inducible expression of immunotoxins in T cells or other effector cells that can localize to tumors.

## Materials and Methods

### Cell lines and cell culture

293FT cells (ThermoFisher #R70007) were cultured in Dulbecco’s Modified Eagle Medium (DMEM) high glucose (25 mM), GlutaMAX, sodium pyruvate (1 mM) (ThermoFisher, #10569010) supplemented with MEM non-essential amino acids (ThermoFisher #11140050), Geneticin (ThermoFisher #10131027), 10% not-heat-inactivated fetal bovine serum (FBS, ThermoFisher #16000044), and penicillin/streptomycin (ThermoFisher #15140122) (293FT media). AsPC1 (ATCC #CRL-1682), Jurkat (ATCC #TIB-152), NALM6 (ATCC #CRL-3273), NCI-H358 (ATCC #CRL-5807), and Ramos RA1 (ATCC #CRL-1596) cells were cultured in Roswell Park Memorial Institute Medium (RPMI)-1640 media (ATCC #30-2001) supplemented with 10% FBS and penicillin/streptomycin. NCI-H358 and NALM6 cells were previously modified to express green fluorescent protein (GFP) and luciferase^49,55,72^. CFPAC-1 cells (ATCC #CRL-1918) were cultured in Iscove’s Modified Dulbecco’s Medium (IMDM; ATCC #30-2005) supplemented with 10% FBS and penicillin/streptomycin. NK-92 cells (ATCC # CRL-2407) were cultured in RPMI-1640 media supplemented with 1% MEM Non-Essential Amino Acids Solution (ThermoFisher #11140050) and IL-2 (Prometheus, NDC #65483-116-07). LV-MAX Viral Production Cells (ThermoFisher #A35347) were cultured in LV-MAX Production Medium (ThermoFisher #A3583401). Primary human CD3+ T cells were isolated using the EasySep Human T Cell Isolation Kit (Stemcell Technologies #17951) from de-identified healthy donor peripheral blood mononuclear cells (PBMCs) and were cultured in RPMI-1640 supplemented with IL-2 (Prometheus, NDC #65483-116-07), IL-7 (BioLegend #581906), 10% FBS, and penicillin/streptomycin (T cell media). Primary human B cells were isolated from de-identified healthy donor PBMC using the EasySep Human B cell Isolation Kit (Stemcell Technologies #17954). Isolated B cells were cultured using the ImmunoCult Human B Cell Expansion Kit (Stemcell Technologies #100-0645) with ImmunoCult-XF B Cell Base Medium (Stemcell Technologies #100-0646) and ImmunoCult-ACF Human B Cell Expansion Supplement (Stemcell Technologies #10974). B-LCLs were generated from de-identified healthy donor PBMCs and were cultured in RPMI-1640 supplemented with 10% FBS and penicillin/streptomycin. Primary human NK cells were isolated from de-identified healthy donor PBMCs using the EasySep Human NK Cell Isolation Kit (Stemcell Technologies #17955). Isolated NK cells were cultured using the ImmunoCult NK Cell Expansion Kit (Stemcell Technologies #100-0711) with ImmunoCult NK Cell Base Medium (Stemcell Technologies #100-0712), ImmunoCult NK Cell Expansion Supplement (Stemcell Technologies #100-0715), and ImmunoCult NK Cell Expansion Coating Material (Stemcell Technologies #100-0714). All cells were cultured at 37°C, 5% CO_2_ unless noted otherwise (e.g., incubations at room temperature, ambient air).

### Synthesis and cloning of immunotoxins

DNA sequences encoding immunotoxin and control proteins were de novo synthesized by GeneArt (ThermoFisher). Constructs were cloned into plasmid vectors using cognate restriction enzymes (New England Biolabs) or NEBuilder HiFi DNA Assembly (NEB #E5520S). Plasmids used were pcDNA3.1 (ThermoFisher #V79020) for DNA transfection, a pUC19 derived vector backbone (Addgene #112021) for CRISPR HDRT^73^, an epHIV7.2 derived vector backbone (gift of C.A. Crane) for lentivirus production^74^, and pCI (Promega #E1731) for mRNA production. Plasmid sequences were confirmed by Sanger sequencing (Azenta Life Sciences). pmaxGFP plasmid (Lonza #V4XP-3032) was used as a transfection control. Sequences are described in Supplementary Table S1 and S2.

### Testing the sensitivity of 293FT cells to ADPRTs

293FT cells were incubated with increasing concentrations of purified, unnicked DT (List Biological Labs #150 or MilliporeSigma #322326) or increasing concentrations of purified PE (MilliporeSigma #P0184). Cell viability was determined using the CellTiter-Glo luminescent cell assay (Promega #G7570) at defined time points. Luminescence was measured as relative light units (RLU) using a GloMAX luminescence microplate reader (Promega), analyzing wells with only cell culture media to correct for background.

### Engineering resistance to ADPRTs in human cell lines

293FT cells were edited with CRISPR/Cas9 to generate specific knockouts of diphthamide biosynthesis genes (*DPH1-7*) or to introduce an *EEF2* p.G717 mutation using homology-directed repair (HDR). S.p. Cas9 crRNAs (IDT, custom synthesis) with specificity for one of the *DPH1-DPH7* gene loci were reconstituted in nuclease-free duplex buffer (IDT #11-01-03-01) and complexed with equimolar ratio of Alt-R CRISPR-Cas9 tracrRNA (IDT #1072534). sgRNA complexes for each *DPH1-7* or *EEF2* gene were incubated with Alt-R S.p. Cas9 Nuclease V3 (IDT #1081058) at a 5:1 molar ratio for 15 minutes at room temperature (RT) to allow for Cas9 ribonucleoprotein (RNP) complex assembly. For HDR of *EEF2*, repair template DNA was synthesized as an Ultramer DNA Oligonucleotide (IDT, custom synthesis). Repair template was incubated with *EEF2*-directed Cas9 RNPs for 10 minutes at RT. CRISPR/Cas9 guide RNA sequences and dsDNA homology-directed repair templates (HDRTs) used are listed in Supplementary Table S1. Adherent 293FT cells were treated with trypsin-EDTA 0.05% (ThermoFisher #25300054) to generate single-cell suspensions. 293FT cells were pelleted by centrifugation at 90 x*g* for 10 minutes. Pelleted cells were resuspended in SF buffer (Lonza #V4XC-2032) at a concentration of 10×10^6^ cells/mL. 293FT cells (0.2×10^6^ cells) in 20 μL SF buffer were combined with either 5 μL Cas9 RNP for *DPH1-7* or Cas9 RNP with HDRT for *EEF2* and 20 μL transferred to each well of a 16-well electroporation cuvette (Lonza). Cells were electroporated in a 4D-Nucleofector X Unit (Lonza, #AAF-1003X) using pulse code CM-130. Electroporated cells were incubated for 5 minutes at RT, 80 μL pre-warmed 293FT media was directly added to each cuvette well, and cells were rested for another 15 minutes at 37°C, 5% CO_2_. Cells were then transferred to 96-well tissue culture treated plates (Corning #3598) at 20,000 cells/well in pre-warmed 293FT media and allowed to recover for 3 days. Following recovery, edited 293FT cell pools were grown in 293FT media containing 1 μg/mL native DT and 1 μg/mL native PE for selection of resistant cells. A combination of two ADPRTs was used to avoid selection for cell clones that lost or acquired mutations in either toxin’s specific cell surface receptor or a key protein of their distinct intracellular trafficking pathways before reaching cytosolic EEF2. Cell proliferation was measured by quantifying cell confluence via live-cell imaging (Sartorius Incucyte S3). At the end of the selection period, all unedited control 293FT cells had died, showing no recovery during subsequent incubation in media without toxin. Edited and selected 293FT cells that acquired ADPRT resistance were then maintained in 293FT media without native ADPRT.

To engineer resistance in Jurkat T cells, 100 pmol sgRNA-1 targeting *DPH1* (IDT custom synthesis) was combined with Alt-R S.p. Cas9 Nuclease V3 (IDT #1081059) and 64 pmol Alt-R Cas9 electroporation enhancer (IDT #1075916) in a total volume of 2.5 μL to generate RNPs. Jurkat T cells were centrifuged at 90 x*g* for 10 minutes and 1×10^6^ cells resuspended in 20 μL SE Buffer (Lonza #V4XC-1032). RNPs (2.5 μL) were combined with equal volumes of Opti-MEM (ThermoFisher #31985070) and 20 μL of Jurkat T cells. Cells were electroporated in a 4D-Nucleofector X Unit using pulse code CL-120. Following electroporation, 80 μL pre-warmed media was added directly to each cuvette well, and the cells were incubated for 30 minutes at 37°C, 5% CO_2_. After recovery, cells were further diluted in pre-warmed media to 10^6^ cells/mL and incubated at 37°C, 5% CO_2_. After 8 days of culture, Jurkat T cells were selected for resistance using 2 μg/mL DT (MilliporeSigma #322326).

### Engineering resistance to ADPRTs in primary human T cells

Analogous to engineering resistance in 293FT cells, we employed an adapted CRISPR/Cas9 editing approach to engineer resistance in primary human T cells. For this, primary human T cells were isolated from healthy donor PBMCs using magnetic bead negative selection (Stemcell Technologies #17951). CD3+ T cells were activated using Dynabeads Human T-Activator CD3/CD28 (ThermoFisher #11131D) at a 1:1 bead:cell ratio in T cell media. At 48 hours of stimulation, Dynabeads were removed and activated T cells were edited using CRISPR/Cas9 (Supplementary Table S1). S.p. Cas9 crRNA (IDT custom synthesis) targeting *DPH1-4* or *EEF2*, as used for 293FT cells, was complexed with Alt-R CRISPR-Cas9 tracrRNA (IDT #1072534) at an equimolar ratio. Complexed sgRNA (100 pmol/10^6^ cells) was incubated with Alt-R S.p. Cas9 Nuclease V3 (50 pmol/10^6^ cells, IDT #1081058) at a 2:1 molar ratio for 15 minutes at RT. For T cell experiments with additional furin knockout, RNPs were produced with guide RNA for the *FURIN* locus and were mixed with DPH-targeting RNP. For homology-directed repair of *EEF2*, 50 pmol of HDRT DNA (IDT custom synthesis) was incubated with *EEF2*-directed Cas9 RNP for 10 minutes at RT. T cells were pelleted by centrifugation at 90 x*g* for 10 minutes and resuspended in Buffer P3 (Lonza #V4XP-3032) at a concentration of 50×10^6^ cells/mL. T cell suspension (1×10^6^ T cells in 20 μL) was combined with Cas9 RNP (for *DPH* knockout) or Cas9 RNP with HDRT (for *EEF2* mutation). The T cell and RNP mixture was transferred to a 16-well electroporation cuvette (Lonza) at 20 μL/well. T cells were electroporated in a 4D-Nucleofector X Unit (Lonza) using pulse code EH-115. Electroporated cells were rested for 5 minutes at room temperature/ambient air, 80 μL pre-warmed T cell media without IL-2 or IL-7 was directly added to each cuvette well, and cells incubated for another 15 minutes at 37°C, 5% CO_2_. Electroporated T cells were then resuspended in 2 mL of T cell media containing IL-2 or IL-7 and allowed to recover by incubation in 12-well tissue culture-treated plates at 37°C, 5% CO_2_. After 3 days, edited and mock-edited (sgRNA control) T cells were resuspended in T cell media containing 1 μg/mL native DT (MilliporeSigma #322326) or no toxin. Cells were seeded in replicate plates or propagated in tissue culture flasks for ongoing selection. T cell proliferation in the presence of DT was quantified using the CellTiter-Glo assay (Promega #G7570) at each day after the start of selection using one replicate plate for each time point. T cell media was replaced in cell culture flasks every 3 days to replenish cytokines and DT.

### Testing viability of ADPRT-resistant 293FT cells

To evaluate the resistance of engineered 293FT cells to ADPRTs, cells were seeded in 293FT media containing 1 μg/mL of native DT (MilliporeSigma #322326), previously established to be lethal to unedited 293FT cells. Cell growth was quantified by longitudinally monitoring the confluence of 293FT cells using live-cell imaging (Sartorius Incucyte S3). To test the susceptibility of engineered 293FT cells to TNFα, cells were seeded in plates in 293FT media containing 1 μg/mL DT, 125 nM Incucyte Cytotox Red Dye (Sartorius #4632), and increasing concentrations of human recombinant TNFα (Cell Signaling Technology #16769). Cell death was quantified over time using live-cell imaging (Sartorius Incucyte S3).

### DNA transfection of 293FT cells with immunotoxin constructs

Wild-type and resistance-engineered 293FT cells were seeded in 96-well tissue culture plates at 20,000 cells/well. Upon reaching 40-50% confluence the next day, cells were transfected using Lipofectamine 3000 reagent (ThermoFisher #L3000001). For this, immunotoxin-encoding or control plasmid DNA (100 ng/well) was diluted in P3000 reagent and Opti-MEM (ThermoFisher #31985062), Lipofectamine reagent was diluted in Opti-MEM, and 10 μL of transfection mixture (1:1 v/v diluted DNA/P3000 and diluted Lipofectamine) was added to cells growing in 200 μL 293FT media. Constructs used for transfection are described in Supplementary Table S2; pCI vector encoding GFP was used as a transfection control, and vector encoding cl13-hUCHT1v9LHLH^55^ was used as a control for anti-FLAG (DYKDDDDK) tag staining. At 48 hours, transfected 293FT cells were analyzed by flow cytometry to evaluate intracellular expression of immunotoxin or control proteins.

### Culture of NCI-H358 target cells with conditioned media

Engineered 293FT cells were first transfected with DNA constructs encoding immunotoxin or control proteins. For this, cells were seeded in 96-well tissue culture plates at 20,000 cells/well. Upon reaching 40-50% confluence the next day, cells were transfected using Lipofectamine 3000 reagent (ThermoFisher #L3000001). Immunotoxin-encoding or control plasmid DNA (100 ng/well) was diluted in P3000 reagent and Opti-MEM (ThermoFisher #31985062), Lipofectamine reagent was diluted in Opti-MEM, and 10 μL of transfection mixture (1:1 v/v diluted DNA/P3000 and diluted Lipofectamine) was added to cells growing in 200 μL 293FT media. Next, 293FT cells were washed and then seeded in the insert chamber of a 24-well transwell culture plate (Corning #3450) at 4×10^4^ cells/well in equal volume 293FT and RPMI-1640 media without Geneticin. NCI-H358 target cells were seeded at 10^4^ cells/well in the lower chamber. NCI-H358 cells were imaged (Sartorius Incucyte S3) by temporarily removing the transwell insert, imaging the lower cell chambers, and then re-inserting the transwell plate for continued co-culture.

### Transwell co-culture of NCI-H358 target cells with effector cells

293FT cells were transfected with DNA constructs encoding immunotoxin or control proteins using Lipofectamine 3000 (ThermoFisher #L3000001), washed, and then seeded in the insert chamber of a 24-well transwell culture plate (Corning #3450) at 4×10^4^ cells per well in equal volume 293FT and RPMI-1640 media without Geneticin. NCI-H358 target cells were seeded at 10^4^ cells per well in the lower chamber. NCI-H358 cells were imaged (Sartorius Incucyte S3) by temporarily removing the transwell insert, imaging the lower cell chambers, and then re-inserting the transwell plate for continued co-culture. NCI-H358 target cells were additionally quantified using the Steady-Glo assay (Promega #E2510). For experiments with primary human T cells, primary human B cells, B-LCL, NALM6, and NK-92, co-cultures were similarly established with 10^5^ engineered effector cells per well.

### Mixed cell co-culture of NCI-H358 target cells with 293FT effector cells

GFP-negative 293FT cells engineered for resistance with Cas9 RNP targeting *DPH1* (DPH1 gRNA-1), *DPH2* (DPH2 gRNA-1), *DPH4* (DPH4 gRNA-2), or *EEF2* (EEF2 gRNA-2 with DNA repair template to introduce p.G717R) were transfected with DNA constructs encoding immunotoxin or control proteins as described above, harvested, and washed. 293FT cells (1.5×10^4^ cells/well) were then added to NCI-H358 target cells (seeded at 5×10^3^ cells/well) in 96-well plates. GFP-positive NCI-H358 cells were quantified longitudinally by live-cell imaging (Sartorius Incucyte S3). 293FT cells that were not transfected served as a negative control.

### Testing viability of ADPRT-resistant primary human T cells

Primary human CD3+ T cells were isolated from PBMCs and *DPH1* was disrupted by CRISPR/Cas9 HDR together with sgRNA-1. For this experiment, 10^6^ primary human T cells were resuspended in 20 μL P2 Buffer or P3 Buffer (Lonza #V4XP-2032 or Lonza #V4XP-3032). Cells were electroporated in a 4D-Nucleofector X Unit (Lonza) using pulse codes EH-100 or EH-111. After 5 days of culture, T cells were activated using Dynabeads Human T-Activator CD3/CD28 (ThermoFisher #11131D) at a ratio of 1:1 beads:cells. At 3 days post-activation, beads were removed using a magnet and T cells resuspended in T cell media containing 2 μg/mL DT. Cells were incubated at 37°C, 5% CO_2_ and imaged using an Incucyte SX5 (Sartorius). T cells were quantified by total phase area per well.

### Functional testing of ADPRT-resistant primary human T cells

For functional assays of ADPRT-resistant T cells, CD3+ primary human T cells were edited using CRISPR/Cas9 and selected in T cell media containing DT for a total of 10 days. Mock-edited CD3+ primary human T cells expanded in T cell media without DT served as controls. Expanded T cells were labeled with CellTrace Violet (ThermoFisher #C34557) and then seeded at 5×10^4^ cells/well in T cell media in 96-well tissue culture plates. GFP+ NALM6 target cells expressing CD19 or edited to knockout CD19 were added at 2.5×10^4^ cells/well. A CD19xCD3 bispecific T cell-engaging antibody (InvivoGen #bimab-hcd19cd3-01) was added to the wells at final concentrations of 0-50 ng/mL. For one plate, 100 μL of cell culture supernatant was harvested after 1 day, centrifuged at 4000 x*g* to remove cell debris, and stored at -80°C for cytokine assays. Cells were incubated for 7 days. At the end of the experiment, cells were pelleted by centrifugation at 350 x*g* for 5 minutes. Co-culture supernatants were harvested as above and frozen at -80°C for cytokine assays. Pelleted cells were subjected to flow cytometry to determine T cell proliferation and cytotoxicity. To do so, cells were stained with LIVE/DEAD Fixable Near-IR (ThermoFisher #L34976), TruStain FcX (BioLegend #422301), and APC anti-human CD3 (BioLegend #981012). Data were acquired using an Intellicyt iQue Screener PLUS (Sartorius) and analyzed using FlowJo v9/10 (BD Life Sciences). All experimental data were analyzed by gating on single, live cells. NALM6 cells were identified as GFP+ and CD3-. T cells were identified as CD3+ and GFP-. Expansion indices were calculated by generating fit curves for CellTrace Violet dilution using the FlowJo v9/10 proliferation platform. For cytokine analyses, supernatants were analyzed using the iQue Qpanel T Helper 6-plex kit (Sartorius #90502), with the modification of diluting supernatants 1:4 in assay buffer.

### Mixed cell co-culture of target cells with Jurkat T cells

Target cells (10^4^ cells/well) were seeded in HTS Transwell polycarbonate membrane (0.4 μm pore) plates (Corning #3391). Jurkat T cells (0.5×10^6^ cells/well) were centrifuged at 90 x*g* for 10 minutes and resuspended in 20 μL SE Buffer (Lonza #V4XC-1032). Plasmid (1 μg in 1 μL) was added to the cells, and 20 μL of the cell mixture was transferred to the electroporation cuvette (Lonza). Jurkat T cells were electroporated in a 4D-Nucleofector X Unit (Lonza) using pulse code CL-120. Electroporated cells were rested for 5 minutes, 80 μL pre-warmed media was directly added to each cuvette well, and the cells were incubated for 30 minutes at 37°C, 5% CO_2_. Cells were diluted to a final density of 0.5×10^6^ cells/mL in media and propagated at 37°C, 5% CO_2_. The following day, Jurkat T cells were added to transwell inserts (5×10^4^ cells/well). An equivalent volume of media without DT or media supplemented with DT (final concentration of 2 μg/mL) was added into the transwell insert of control wells. Cells were imaged using an Incucyte SX5 (Sartorius) and cell density measured as total phase area of cells per well. Jurkat T cells were removed with the transwell insert after 48 hours of co-culture.

### Production of lentiviral particles

LV-MAX Viral Production Cells were CRISPR edited for knockout of *DPH1* and then selected for ADPRT resistance by passaging in media supplemented with exogenous DT as described above for 293FT cells. Resistant LV-MAX cells were transfected and cultured using the LV-MAX Lentiviral Production System (ThermoFisher #A35684). Supernatant was collected and concentrated using Lenti-X Concentrator (Takara #631231). Lentiviral particle concentration was measured using the QuickTiter Lentivirus Titer Kit (Cell Biolabs #VPK-107).

### Lentiviral transduction

Primary human T cells were engineered in a two-step process in which *DPH1* was first knocked out with CRISPR and cells were subsequently transduced with immunotoxin construct lentiviral particles^75–78^. Primary human T cells were isolated from de-identified healthy donor PBMCs and cultured overnight. The next day, unactivated T cells were subjected to CRISPR editing of *DPH1* as described above. Two days later, T cells were activated with a 3:1 bead:cell ratio of Dynabeads Human T-Activator CD3/CD28 (ThermoFisher #1132D). After approximately 24 hours of activation, lentiviral particles were added to T cells. Media was changed two days later. Cells were then used for downstream applications. For transduction of 293FT, lentiviral particles were added to cells growing in log phase at approximately 50% confluency. Cells were passaged one day later.

### CRISPR homology-directed repair knockin

Homology-directed repair templates (HDRT) were generated by PCR amplification of constructs from plasmids using Q5 Hot Start High-Fidelity 2X Master Mix (NEB #M0494S). HDRT were purified after PCR with AMPure XP (Beckman Coulter #A63881). For T cell knockin, primary human T cells were isolated from de-identified healthy donor PBMCs. After overnight culture, unactivated T cells were CRISPR edited for *DPH1* as described above. Media was changed at 2 days and 4 days post-CRISPR editing. On day 5, T cells were activated with a 1:1 bead:cell ratio of Dynabeads Human T-Activator CD3/CD28 (ThermoFisher #1132D). After approximately 24 hours, T cell knockin was performed using CRISPR as described above with the following modifications: a guide RNA for the beta-2 microglobulin locus was used to create RNPs and these RNPs were then complexed with 0.5 μg HDRT for 5 minutes at room temperature^73^. T cells were then used for downstream applications. For constructs using Cre recombination, HDRTs were designed such that loxP sites flanked immunotoxin constructs in an inverted orientation. T cells were edited in a single round of CRISPR using HDRTs and guide RNA targeting the *DPH1* locus. Five days post-editing, T cells were treated with TAT-CRE recombinase (MilliporeSigma #SCR508) to recombine the construct insert into a direction compatible with transcription.

### mRNA transfection

mRNA was generated by linearization of plasmids with cognate restriction enzymes (NEB), in vitro transcription with capping and tailing using the T7 mScript Standard mRNA production system (CELLSCRIPT C-#MSC11610 and #C-MSC100625), and purification using the MEGAclear Transcription Clean-Up Kit (ThermoFisher # AM1908). Alternatively, mRNA was synthesized in custom production (ProMab Biotechnologies). mRNA lipid nanoparticles (mRNA-LNP) were also synthesized in custom production (ProMab Biotechnologies). Primary human T cells were isolated from PBMCs and activated using a 1:1 bead:cell ratio of Dynabeads Human T-Activator CD3/CD28 (ThermoFisher #1132D). After approximately 48 hours of activation, T cells were edited with CRISPR for *DPH1* knockout as described above. Approximately 14 days after activation, 2.5×10^6^ T cells mixed with 10 μg RNA in 100 μl Opti-MEM (ThermoFisher #31985070) were electroporated with 200 V for 16 ms (BTX #ECM 2001). For mRNA-LNP transfection, mRNA-LNPs were added to T cells and gently mixed. 293FT and NCI-H358 cells were transfected using Lipofectamine MessengerMAX (ThermoFisher #LMRNA00) or mRNA-LNP.

### Flow cytometric evaluation

For flow cytometry evaluation of intracellular immunotoxin constructs, 293FT cells were treated with trypsin-EDTA 0.05% (ThermoFisher #25300054) to obtain a single-cell suspension. T cells were mixed to achieve single cell suspension. Cells were then stained with LIVE/DEAD Fixable Near-IR (ThermoFisher #L34976). Cells were next fixed and permeabilized using Cyto-Fast Fix/Perm buffers (BioLegend #426803) and stained with anti-FLAG-tag antibody (BioLegend #637322). For evaluation of expression of a linked CAR construct, staining was performed with a cognate tetramer consisting of KRAS p.G12V neoantigen peptide^55^ VVGAVGVGK (Peptide 2.0 custom synthesis) complexed with HLA-A*03 (Fred Hutchinson Cancer Center Immune Monitoring Services custom tetramer production). Alternatively, cells were stained with biotinylated recombinant protein L (ThermoFisher #29997) followed by secondary staining with anti-biotin antibody (Miltenyi #130-113-29). Data were acquired using an Intellicyt iQue Screener PLUS (Sartorius) instrument and analyzed using FlowJo v9/10 (BD). Experimental data were analyzed by gating on single, live cells.

### Generation of ADPRT resistant primary human B cells

Primary human B cells were isolated from de-identified healthy donor PBMCs using the EasySep Human B Cell Isolation Kit (Stemcell Technologies #17954). Isolated B cells were cultured using the ImmunoCult Human B Cell Expansion Kit (Stemcell Technologies #100-0645) with ImmunoCult-XF B Cell Base Medium (Stemcell Technologies #100-0646) and ImmunoCult-ACF Human B Cell Expansion Supplement (Stemcell Technologies #10974). After approximately 48 hours, B cells were CRISPR edited for *DPH1* knockout as described above for T cells^79,80^. Buffer P3 and pulse code EH-115 were used. B cells were passaged 3 days later. After 2 further days of culture, B cells were transfected with immunotoxin construct mRNA-LNP and used for further experiments.

### Generation of ADPRT resistant B-LCL, NALM6, and NK-92 cells

B-LCLs were generated from healthy donor PBMCs by Epstein-Barr virus transformation (Johns Hopkins Genetic Resources Core Facility). B-LCLs were edited with CRISPR for *DPH1* knockout as described for T cells using Buffer SF and pulse code DN-100. NALM6 cells were CRISPR edited using Buffer SF with pulse code CV-104. NK-92 cells were edited in Buffer SE with pulse code CA-137. After editing, ADPRT resistant cells were selected by passaging in media supplemented with DT before use in downstream experiments. For immunotoxin construct testing, cells were electroporated with expression plasmids as described above for Jurkat cells using Buffer SF with pulse code DN-100 for B-LCL, Buffer SF with pulse code CV-104 for NALM6, and Buffer SE with pulse code CA-137 for NK-92.

### Generation of ADPRT resistant primary human NK cells

Primary human NK cells were isolated from de-identified healthy donor PBMCs using the EasySep Human NK Cell Isolation Kit (Stemcell Technologies #17955). After isolation, NK cells were cultured with the ImmunoCult NK Cell Expansion Kit (Stemcell Technologies #100-0711) using ImmunoCult NK Cell Base Medium (Stemcell Technologies #100-0712), ImmunoCult NK Cell Expansion Supplement (Stemcell Technologies #100-0715), and ImmunoCult NK Cell Expansion Coating Material (Stemcell Technologies #100-0714). NK cells were CRISPR edited for *DPH1* knockout after 5 days of growth as for T cells described above using Buffer P3 and pulse code EH-115^79,80^. For expression of immunotoxin constructs, edited NK cells were transfected with mRNA-LNP.

### Statistical analysis

Data are shown as individual points or mean ± SEM. Statistical tests were calculated using Prism 10.6.1(GraphPad) or R 4.4. Tests are described in figure legends.

## Acknowledgements

We thank Ali Dbouk, Kathleen Helwig, Alisha Mills, and Leslie Wang for laboratory support and Elizabeth Cook for illustrations. We thank Tolulope Awosika, Kathy Gabrielson, Jiaxin Ge, Yang Li, Jin Liu, Nikita Marcou, Ashley Cook Morgan, Brock Moritz, Tushar Nichakawade, Surojit Sur, Sze-Kat Tan, Joshua Urban, Katherine Wright, Nicolas Wyhs, and Yuan Xia for insightful discussions.

## Funding

Alex’s Lemonade Stand Foundation (C.B.), American Society of Hematology Scholar Award (S.P.), Bloomberg∼Kimmel Institute for Cancer Immunotherapy (K.W.K., D.M.P., B.V., S.Z.), Burroughs Wellcome Career Award for Medical Scientists (C.B.), Commonwealth Fund (C.B., N.P.), Conrad Hilton Foundation (K.W.K., N.P., B.V.), Cupid Foundation (M.F.K.), Jennison Family Funds (C.B.), Jerome Greene Foundation Scholar and Discovery Awards (M.F.K.), Leukemia and Lymphoma Society Translation Research Program award (S.P.), Lustgarten Foundation for Pancreatic Cancer Research (B.V.), NIH grant CA06973 (K.W.K., N.P., B.V.), NIH grant K08CA270403 (S.P.), NIH grant P30CA006973, NIH grant 1R21AI176764-01 (M.F.K.), NIH grant R01CA276221 (C.B.), NIH grant R21TR004059 (C.B.), NIH grant RA37CA230400 (C.B.), NIH grant T32AR048522 (M.F.K.), NIH grant T32GM136577 (S.R.D., J.D., B.J.M., A.H.P.), NIH grant U01CA230691 (C.B., N.P.), Reza Khatib Brain Tumor Center (C.B.), Sol Goldman Sequencing Facility at Johns Hopkins (B.V.), Swim Across America Translational Cancer Research Award (S.P.), Thomas M. Hohman Memorial Cancer Research Fund (C.B.), Virginia and D.K. Ludwig Fund for Cancer Research (C.B., K.W.K., N.P., B.V.).

## Data and Materials Availability

All data is available in the manuscript and supplementary materials. Plasmids are available upon request under a material transfer agreement with The Johns Hopkins University, within the constraints of government regulations regarding the sharing of engineered biological toxins.

## Supplementary Materials

Supplementary Table S1: CRISPR sequences.

Supplementary Table S2: Construct sequences.

Supplementary Figure S1: Disruption of diphthamide biosynthesis pathway proteins and selective EEF2 mutations confer resistance to ADPRT.

Supplementary Figure S2: CRISPR/Cas-engineered diphthamide biosynthesis protein deficient and EEF2 p.G717R mutant 293FT cells are resistant to bacterial ADPRTs.

Supplementary Figure S3: Proliferation of diphthamide biosynthesis protein deficient and EEF2 p.G717R mutant T cells in the presence of ADPRT.

Supplementary Figure S4: Proliferation of ADPRT-resistant engineered T cells in the presence of target cells.

Supplementary Figure S5: Sensitivity of engineered cells with DPH or EEF2 mutations to TNFα.

Supplementary Figure S6: Designs of immunotoxin and control constructs tested in experiments.

Supplementary Figure S7: Expression of GFP in different engineered 293FT cells used as a transfection control.

Supplementary Figure S8: NCI-H358 target cell growth is inhibited by conditioned media supernatant from 293FT cells transfected with immunotoxin constructs.

Supplementary Figure S9: Morphological changes of intoxication and cell death in NCI-H358 target cells.

Supplementary Figure S10: NCI-H358 target cell viability following exposure to immunotoxin producing 293FT cells.

Supplementary Figure S11: Treatment of NCI-H358 target cells with engineered 293FT cells expressing immunotoxins.

Supplementary Figure S12: Designs of immunotoxin constructs with additional targeting and catalytic domains.

Supplementary Figure S13: Lack of robust ADPRT expression in primary human T cells.

Supplementary Figure S14: Limitations of functionality of ADPRTs in primary human T cells.

Supplementary Figure S15: Restricted utility of alternative ADPRT constructs in primary human T cells.

Supplementary Figure S16: Limited ADPRT expression in alternative primary human cell types.

Supplementary Video S1: 293FT DPH1.KO mock x H358 co-culture (day 0-4).

Supplementary Video S2: 293FT DPH1.KO tEGF-PE38 x H358 co-culture (day 0-4).

## Notes

### Competing Interest Statement

M.F.K. is a consultant to Allogene, Amgen, Argenx, Atara Biotherapeutics, Bristol Myers Squibb, Mucommune, Revel Pharmaceuticals, Sana Biotechnology, and Sanofi. M.F.K. receives research support from Blackbird Labs and TBD Pharmaceuticals, Inc. S.P. is a consultant to Merck, owns equity in Gilead, and received payment from QVIA and Curio Science. M.F.K., S.P., B.V., K.W.K., D.M.P., and S.Z. are founders of, hold equity in, and are consultants to TBD Pharmaceuticals, Inc. N.P. and C.B. are consultants to TBD Pharmaceuticals, Inc. K.W.K., N.P., and B.V. are founders of Thrive Earlier Detection, an Exact Sciences Company. K.W.K. and N.P. are consultants to Thrive Earlier Detection. B.V., K.W.K., N.P., and S.Z. hold equity in Exact Sciences. B.V., K.W.K., N.P., and S.Z. are founders of or consultants to and own equity in Clasp, Neophore, and Personal Genome Diagnostics. B.V., K.W.K., and N.P. hold equity in Haystack Oncology and CAGE Pharma. N.P. is a consultant to Vidium. B.V. is a consultant to and holds equity in Catalio Capital Management. S.Z. has a research agreement with BioMed Valley Discoveries. C.B. is a consultant to Depuy-Synthes, Bionaut Labs, Haystack Oncology, Galectin Therapeutics, and is a co-founder of OrisDx and Belay Diagnostics. D.M.P. reports grant and patent royalties from Bristol Myers Squibb, a grant from Compugen, stock from Trieza Therapeutics and Dracen Pharmaceuticals, founder equity from Potenza, is a consultant to Aduro Biotech, Amgen, Astra Zeneca, Bayer, DNAtrix, Dynavax Technologies Corporation, Ervaxx, FLX Bio, Rock Springs Capital, Janssen, Merck, Tizona and Immunomic Therapeutics, is on the scientific advisory board of Five Prime Therapeutics, Camden Nexus II, WindMil, and is on the board of directors for Dracen Pharmaceuticals. J.D. is a consultant to Hemogenyx Pharmaceuticals. The companies named above, as well as other companies, have licensed previously described technologies from The Johns Hopkins University. Licenses to these technologies are or will be associated with equity or royalty payments to the inventors as well as to Johns Hopkins University. The Johns Hopkins University has filed provisional patent applications related to technologies described in this paper on which K.W.K., M.F.K., A.P., N.P., B.V., and S.Z. are listed as inventors. The terms of these arrangements are being managed by The Johns Hopkins University according to its conflict-of-interest policies.

