## Supplementary Figures for "Engineering immunotoxin-equipped effector cells and evaluation in primary human immune cells"

#### Supplementary Figure S1

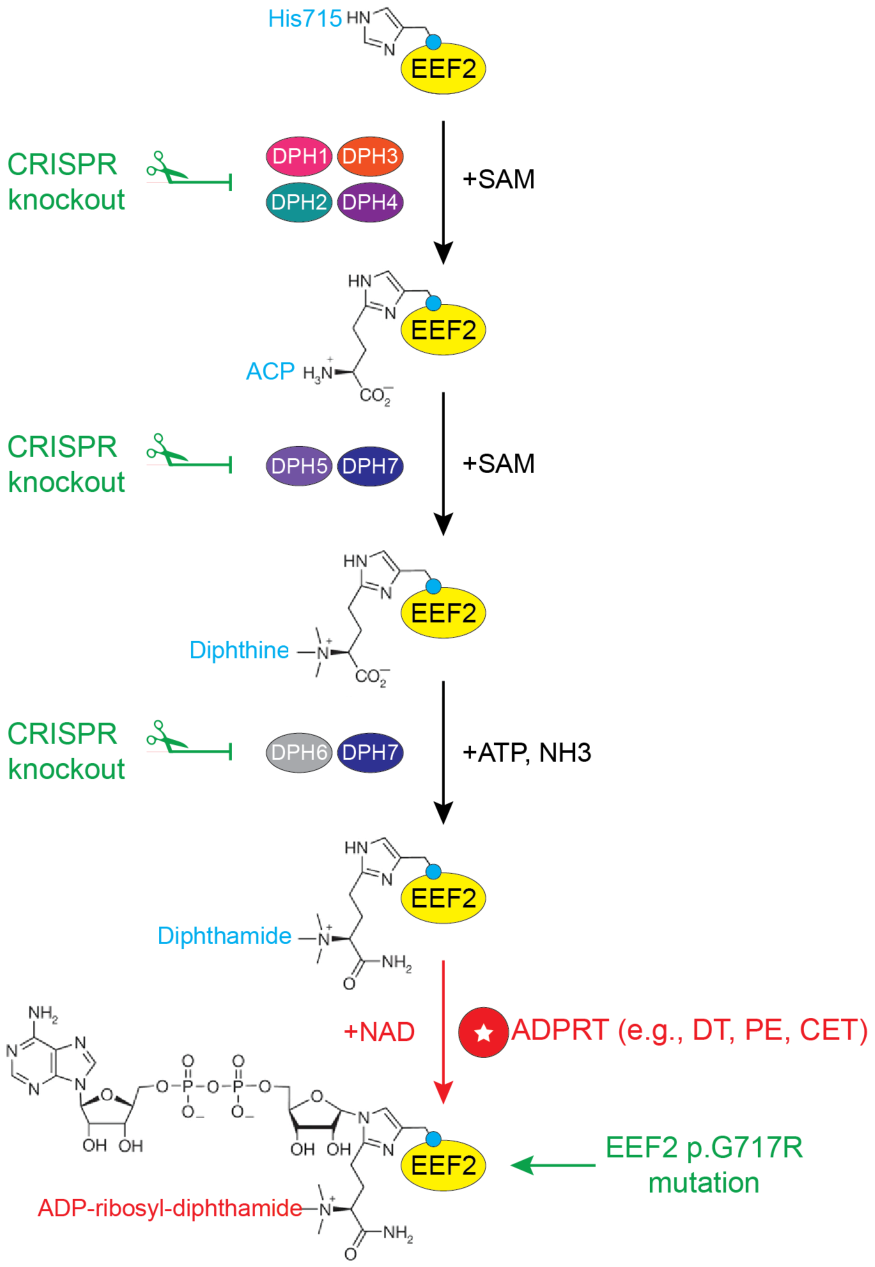


**Supplementary Figure S1: Disruption of diphthamide biosynthesis pathway proteins and selective EEF2 mutations confer resistance to ADPRT.** Diphthamide synthesis is a multistep reaction catalyzed by a group of seven enzymes (DPH1-7), resulting in the posttranslational modification of histidine residue 715 of human EEF2. ADP-ribosylation of the diphthamide residue on EEF2 by bacterial ADPRT, including DT, PE, and CET, results in abrogation of protein synthesis and secondary cell death. Mutation of EEF2 at residue 717 (p.G717R) prevents posttranslational modification of H715 by diphthamide, rendering EEF2 resistant to the action of ADPRTs in generating ADP-ribosyl-diphthamide and thereby halting protein synthesis. ADP: adenosine diphosphate. ADPRT: adenosine diphosphate-ribosylating toxin. CET: *Vibrio cholerae* exotoxin. DT: *Corynebacterium diphtheria* toxin. EEF2: eukaryotic elongation factor 2. PE: *Pseudomonas aeruginosa* exotoxin A.

#### Supplementary Figure S2

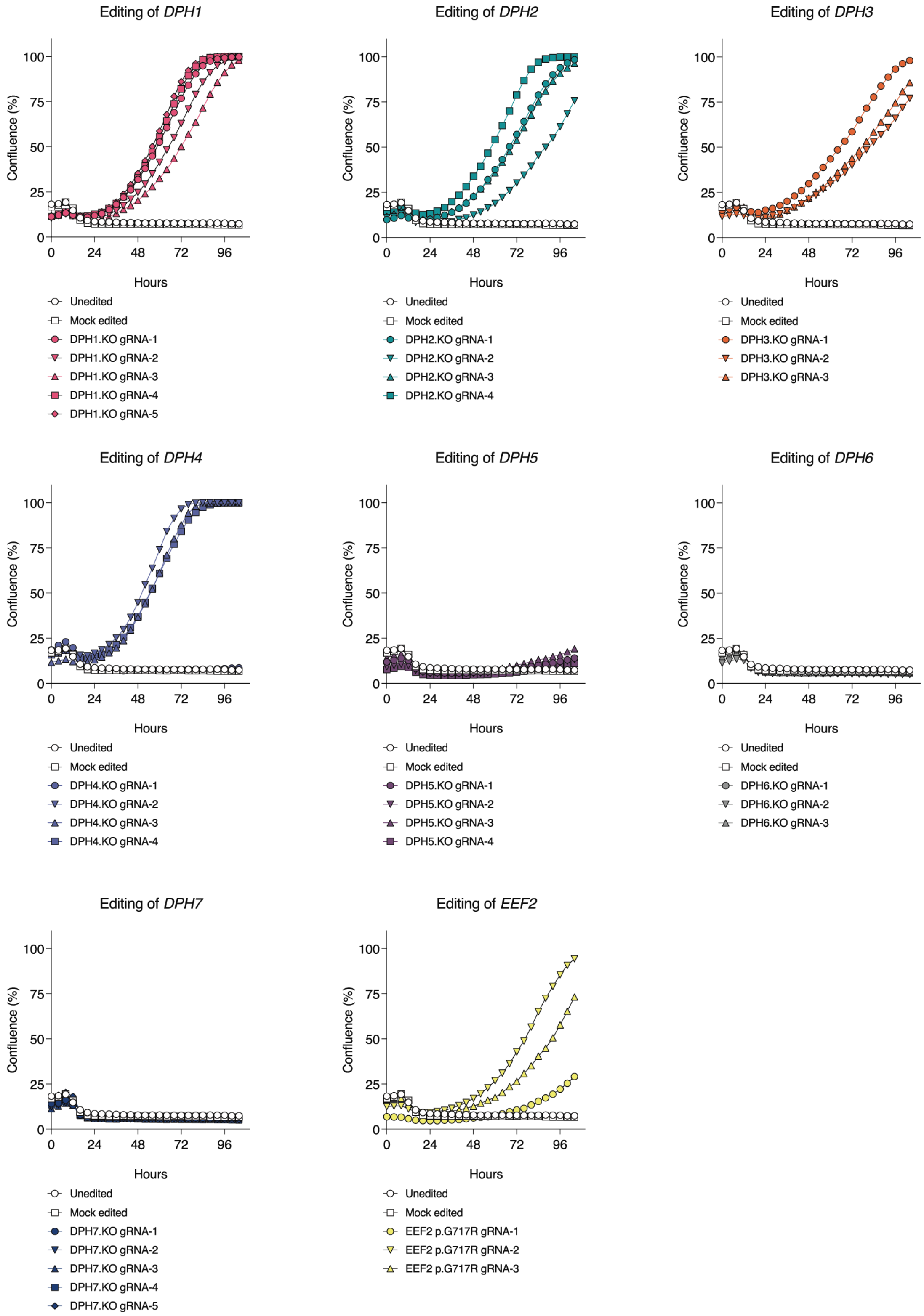

**Supplementary Figure S2: CRISPR/Cas-engineered diphthamide biosynthesis protein deficient and EEF2 p.G717R mutant 293FT cells are resistant to bacterial ADPRTs.** Human 293FT cells were edited using CRISPR/Cas9 to knockout genes of the diphthamide biosynthesis pathway, namely *DPH1*, *DPH2*, *DPH3*, *DPH4*, *DPH5*, *DPH6*, and *DPH7* using up to 5 different guide RNAs (designated gRNA-1 to gRNA-5) per gene. Alternatively, CRISPR/Cas9-mediated homology-directed repair was used to introduce a p.G717R mutation into EEF2, the target of bacterial ADPRTs. Edited 293FT cells were then cultured in the presence of native DT (1000 ng/mL), and their proliferation was quantified over time by measuring confluence using live-cell imaging. ADPRT: adenosine diphosphate ribosylating toxin. DT: *Corynebacterium diphtheria* toxin. EEF2: eukaryotic elongation factor 2.

#### Supplementary Figure S3

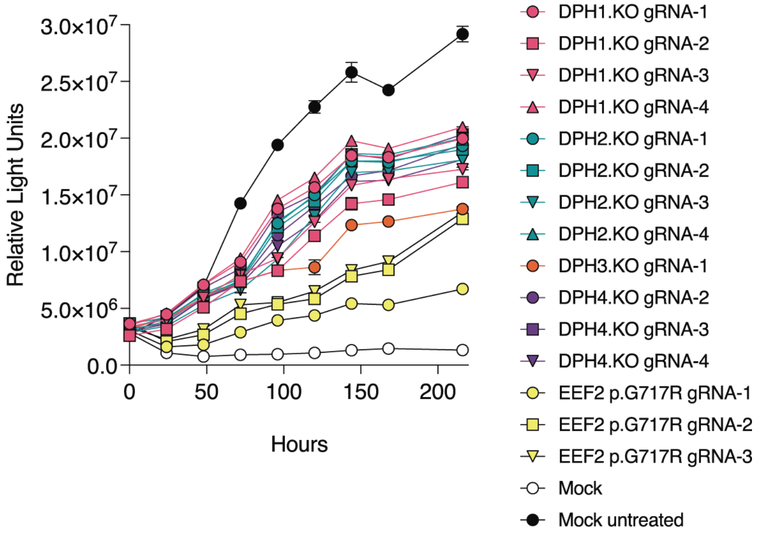


**Supplementary Figure S3: Proliferation of diphthamide biosynthesis protein deficient and EEF2 p.G717R mutant T cells in the presence of ADPRT.** Primary human T cells were edited via CRISPR/Cas9 to knockout *DPH1*, *DPH2*, *DPH3*, or *DPH4* (DPH.KO) or to introduce an EEF2 p.G717R mutation, testing multiple guide RNAs per gene. T cell expansion in the presence of 1 mg/mL native DT was quantified from replicate plates at different time points using the CellTiter-Glo luciferase assay. Mock edited cells were subjected to the CRISPR/Cas9 editing protocol without guide RNA. Untreated cells were cultured in the absence of DT. DT: *Corynebacterium diphtheria* toxin. EEF2: eukaryotic elongation factor 2.

#### Supplementary Figure S4

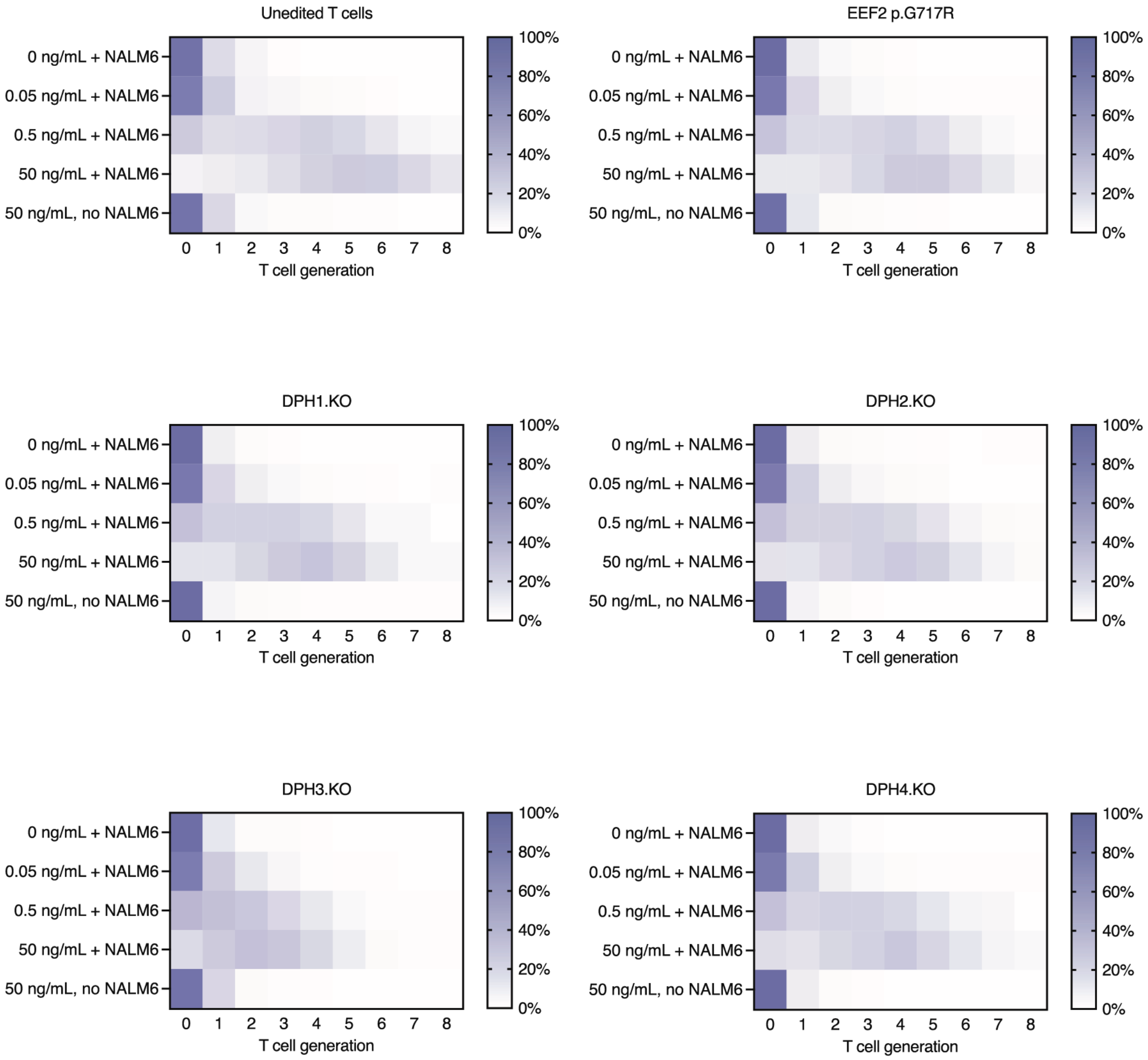


**Supplementary Figure S4: Proliferation of ADPRT-resistant engineered T cells in the presence of target cells.** CRISPR/Cas9 was used to knockout *DPH1*, *DPH2*, *DPH3*, or *DPH4* (DPH.KO) or to introduce an EEF2 p.G717R mutation in primary human T cells. After selection in DT, T cells were co-cultured with CD19+ NALM6 target cells and a CD19xCD3 bispecific T cell engaging antibody at concentrations of 0, 0.05, 0.5, or 50 ng/mL. Percentages of CellTrace Violet labeled T cells in each replication generation as quantified by flow cytometry are shown. DT: *Corynebacterium diphtheria* toxin. EEF2: eukaryotic elongation factor 2.

### Supplementary Figure S5

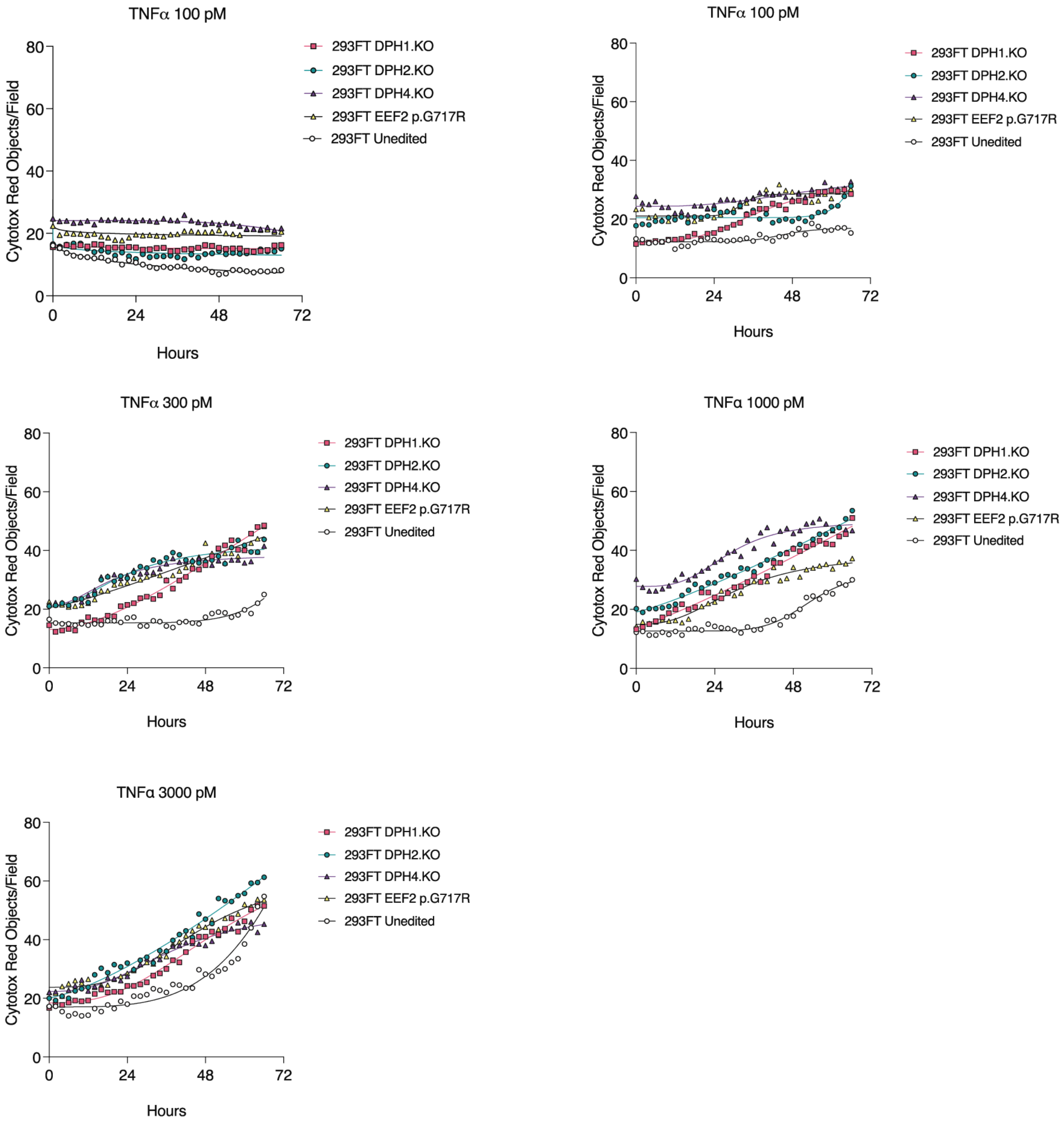


**Supplementary Figure S5: Sensitivity of engineered cells with DPH or EEF2 mutations to TNFα.** 293FT cells edited with CRISPR/Cas9 for *DPH1*, *DPH2*, or *DPH4* knockout (DPH.KO) or EEF2 p.G717R mutation were selected in DT and then cultured in the presence of varying concentrations of exogenous TNFα. Cell death over time was quantified by live cell microscopy using the Cytotox Red marker. DT: *Corynebacterium diphtheria* toxin. EEF2: eukaryotic elongation factor 2. TNFα: tumor necrosis factor-α.

#### Supplementary Figure S6

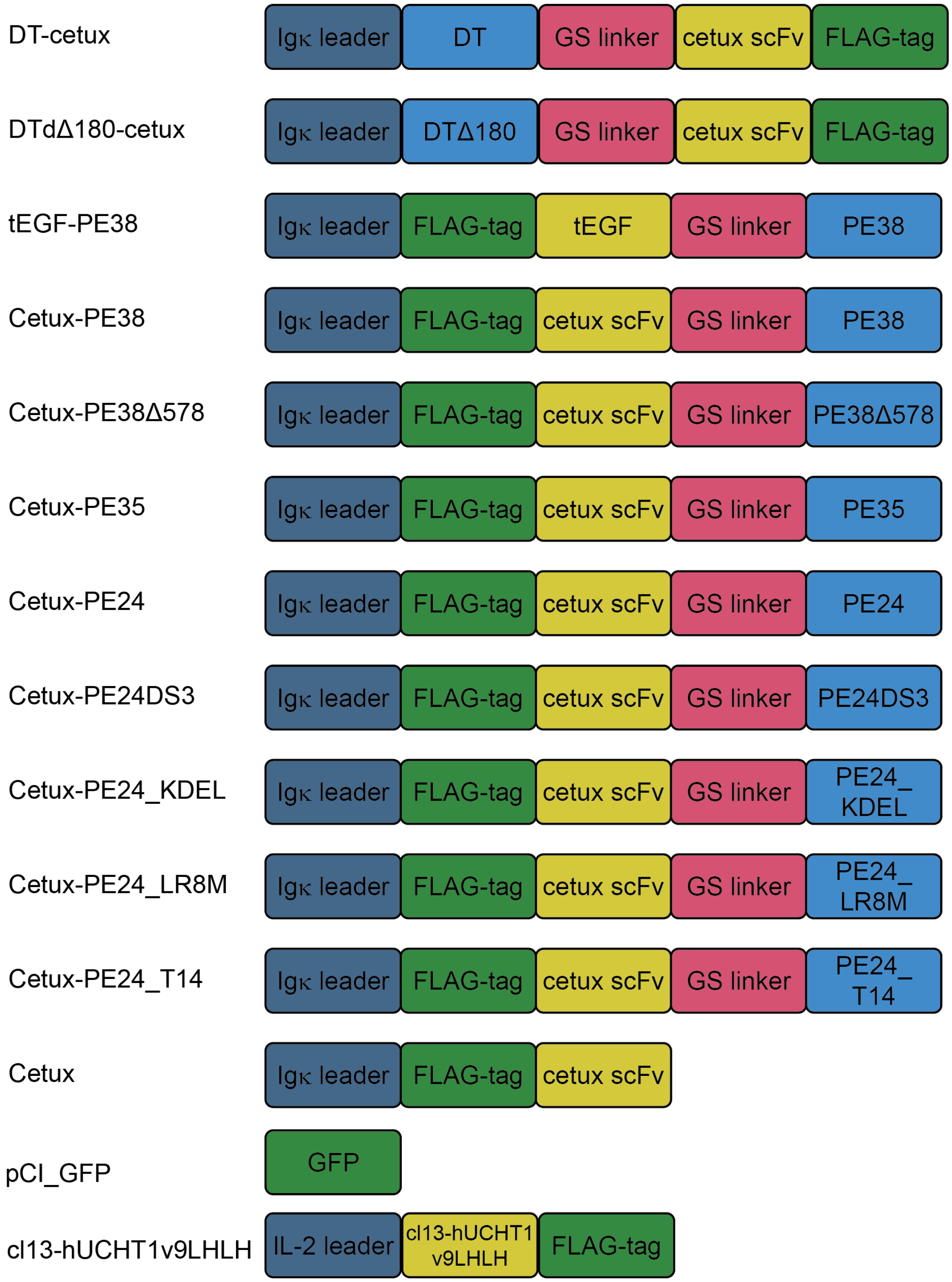


**Supplementary Figure S6: Designs of immunotoxin and control constructs tested in experiments.** Full sequences are described in Supplementary Table S2.

#### Supplementary Figure S7

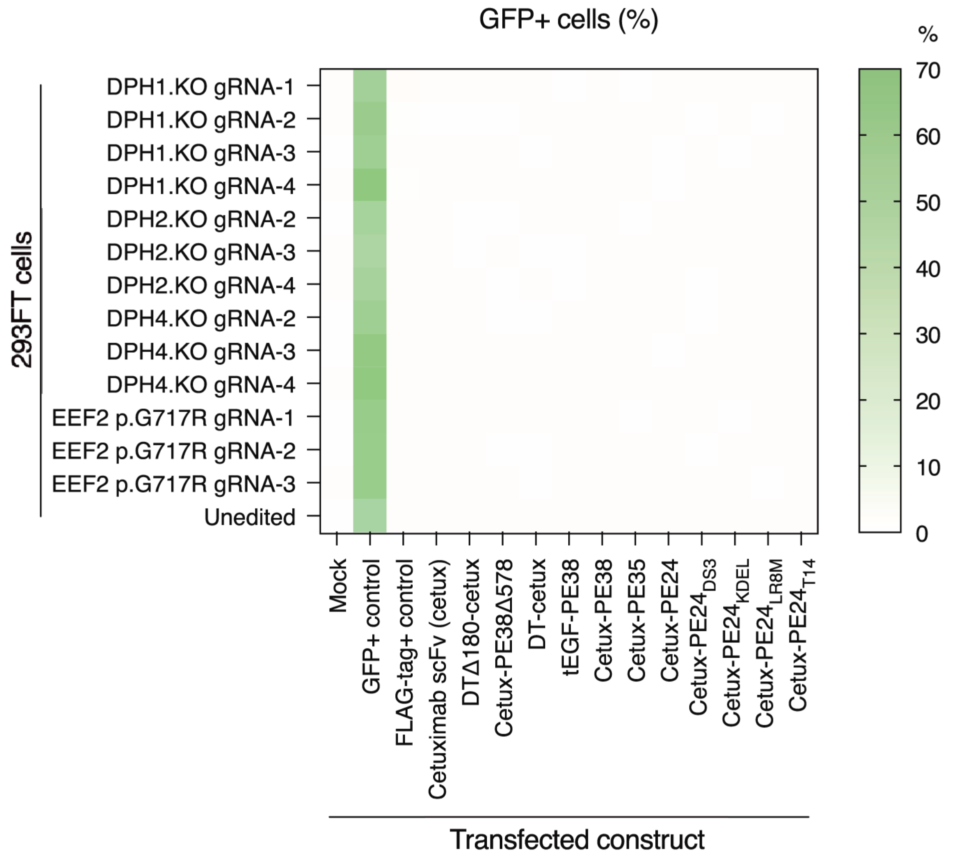


**Supplementary Figure S7: Expression of GFP in different engineered 293FT cells used as a transfection control.** 293FT cells engineered with resistance by CRISPR/Cas9 editing of *DPH* genes (DPH.KO) or introduction of EEF2 p.G717R mutation were transfected for expression of immunotoxin or control constructs. The percentage of live cells expressing GFP was quantified by flow cytometry. EEF2: eukaryotic elongfation factor 2.

#### Supplementary Figure S8

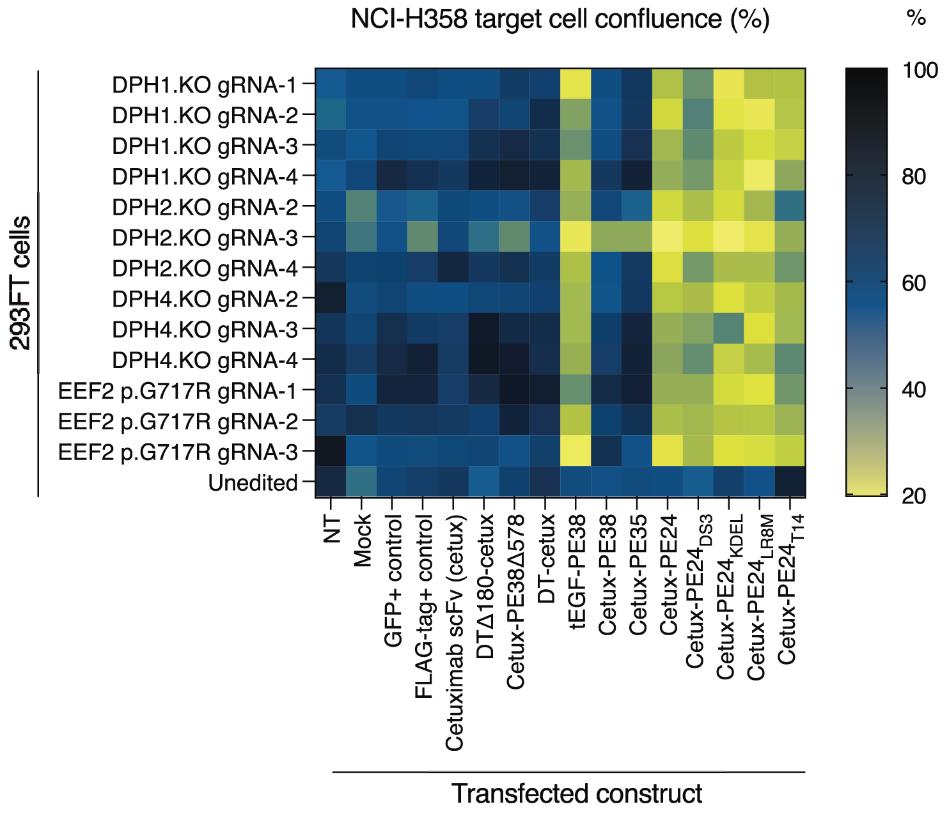


**Supplementary Figure S8: NCI-H358 target cell growth is inhibited by conditioned media supernatant from 293FT cells transfected with immunotoxin constructs.** 293FT cells engineered with resistance by CRISPR/Cas9 editing of *DPH* genes (DPH.KO) or introduction of EEF2 p.G717R mutation were transfected for expression of immunotoxin or control constructs. Conditioned media supernatant from transfected 293FT cells was applied to NCI-H358 target cells. NCI-H358 cell confluence was measured by microscopy after 4 days of supernatant exposure. EEF2: eukaryotic elongation factor 2. NT: non-transfected.

#### Supplementary Figure S9

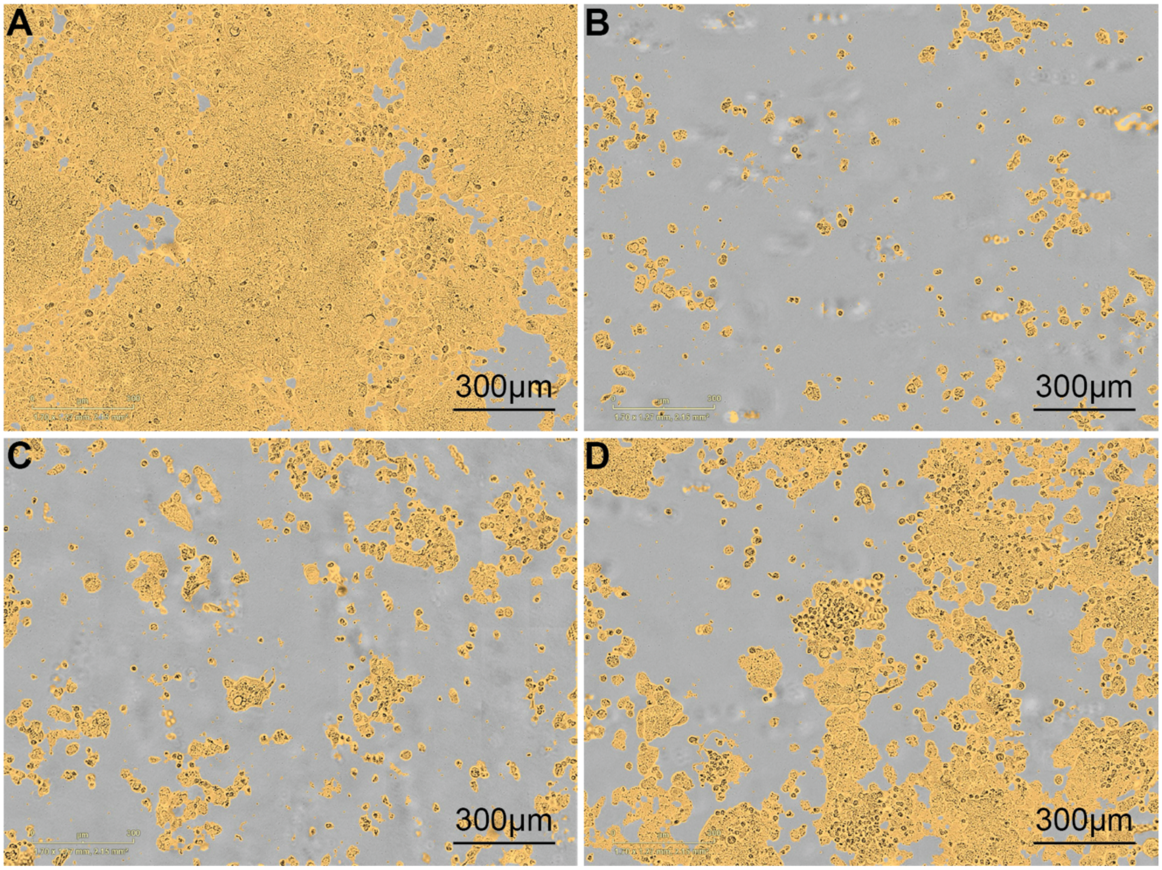


**Supplementary Figure S9: Morphological changes of intoxication and cell death in NCI-H358 target cells.** 293FT cells engineered for resistance with *DPH2* knockout were mock-transfected or transfected with different immunotoxin constructs. (A) Supernatant from non-transfected control 293FT cells, (B) 293FT cells transfected with tEGF-PE38, (C) 293FT cells transfected with anti-EGFR cetuximab scFv-$PE24_{KDEL}$, or (D) 293FT cells transfected with anti-EGFR cetuximab scFv-$PE24_{T14}$ was applied to NCI-H358 target cells. NCI-H358 cell morphology was examined by live cell imaging after 7 days of supernatant exposure. Cells are highlighted with yellow mask.

#### Supplementary Figure S10

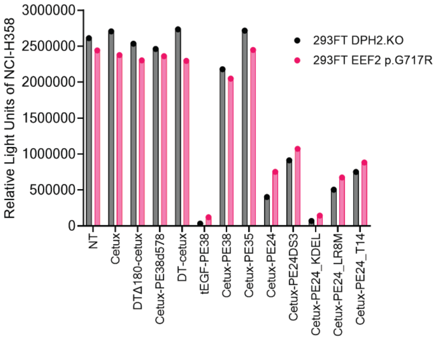


**Supplementary Figure S10: NCI-H358 target cell viability following exposure to immunotoxin producing 293FT cells.** 293FT cells engineered for resistance with *DPH2* knockout (DPH2.KO) or *EEF2* p.G717R mutation were transfected with immunotoxin constructs and then cultured across a transwell barrier insert from NCI-H358 target cells. Viability of NCI-H358 target cells was determined by quantifying luciferase activity (SteadyGlo assay) after 9 days of culture. This endpoint corresponds to the data shown in Fig. 4C and Supplementary Fig. S11A. NT: non-transfected.

#### Supplementary Figure S11

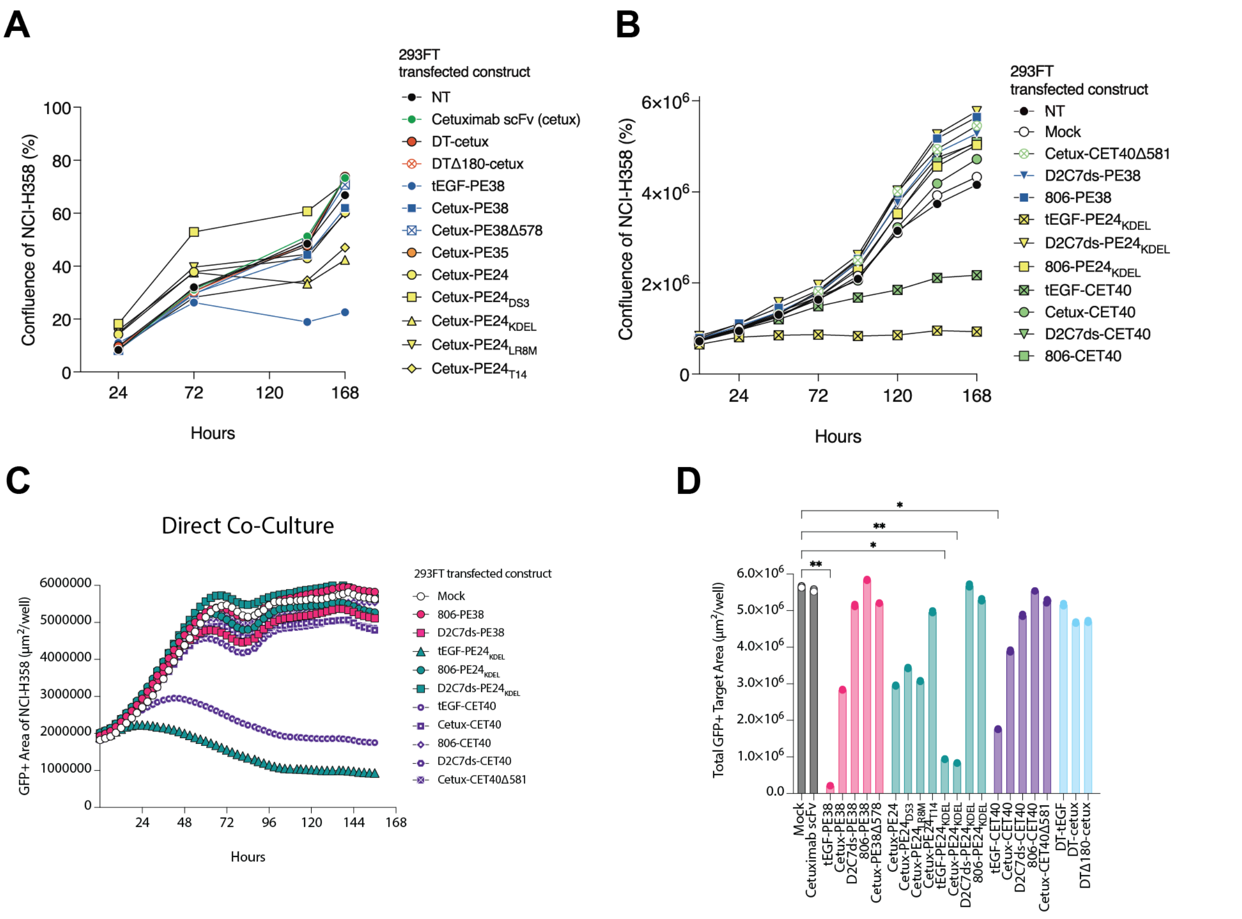


**Supplementary Figure S11: Treatment of NCI-H358 target cells with engineered 293FT cells expressing immunotoxins.** 293FT cells engineered for ADPRT resistance were transfected with different immunotoxin constructs and cultured with EGFR+ NCI-H358 target cells. (A) 293FT cells engineered for resistance with EEF2 p.G717R mutation were transfected with immunotoxin constructs as used in Fig. 4C and co-cultured across a transwell insert with NCI-H358 cells. Confluence of NCI-H358 cells was quantified using live cell microscopy. (B) 293FT cells engineered for resistance were transfected with CET40-based immunotoxin constructs and those with alternative binding moeities, followed by co-culture across a transwell insert with NCI-H358 cells. Confluence of NCI-H358 cells was quantified using live cell microscopy. (C) 293FT cells engineered for resistance were transfected with immunotoxin constructs used in (B) and directly co-cultured with NCI-H358 target cells cells. GFP+ NCI-H358 cells were quantified longitudinally using live cell microscopy. (D) Quantification of NCI-H358 cells in direct co-culture with all resistance-engineered, transfected 293FT cells at experiment endpoints shown in Fig.4F and Supplementary Fig. S11C. *: $p<0.05$; **: $p<0.01$ by Kruskal-Wallis test with Dunn's test.

#### Supplementary Figure S12

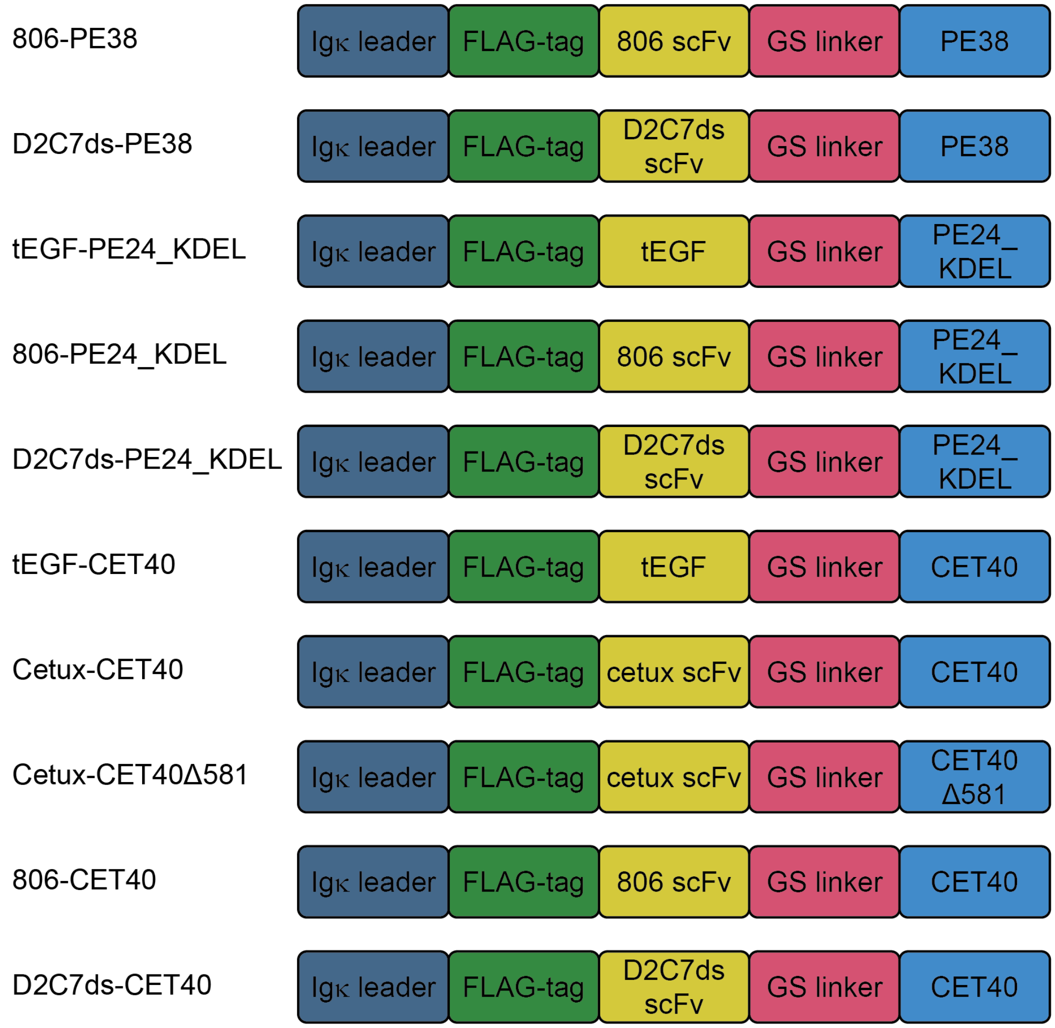


**Supplementary Figure S12: Designs of immunotoxin constructs with additional targeting and catalytic domains.** Full sequences are described in Supplementary Table S2.

#### Supplementary Figure S13

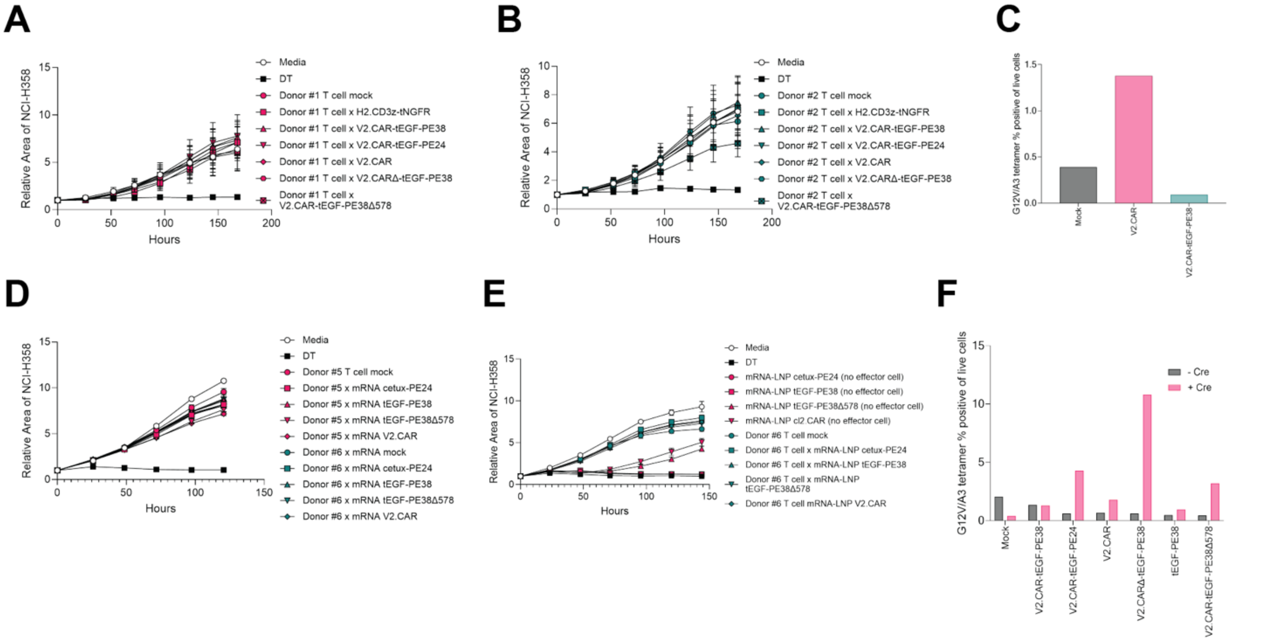


**Supplementary Figure S13: Lack of robust ADPRT expression in primary human T cells.** (A, B) Primary human T cells derived from 2 separate donors were engineered for resistance to ADPRTs using CRISPR and then were transduced with immunotoxin or control constructs. Cells were then co-cultured across a transwell barrier from NCI-H358 target cells. Cell area across time was quantified. (C) Primary human T cells were engineered for resistance to ADPRTs using CRISPR and then engineered in a second CRISPR round for knockin of control or immunotoxin constructs. Expression of a linked CAR functioning as an expression marker was assayed by flow cytometry staining with a cognate tetramer. (D) Primary human T cells were electroporated for transfection of mRNA constructs for expression of control or ADPRT constructs. Cells were then cultured across a transwell barrier from NCI-H358 target cells and target cell area was measured across time. (E) NCI-H358 target cells were either treated directly with immunotoxin mRNA-LNP constructs or were cultured across a transwell barrier from primary human T cells transfected with immunotoxin mRNA-LNP. H358 target cell area across time was measured with live cell microscopy. (F) Constructs containing Cre recombinase sites together with ADPRT modules in an inverted orientation were knocked into the *DPH1* locus in primary human T cells. These cells were then treated with exogenous Cre recombinase to re-orient the ADPRT sequences into an orientation compatible with transcription. Expression was assayed by flow cytometry staining with a cognate tetramer for a linked CAR construct marker.

#### Supplementary Figure S14

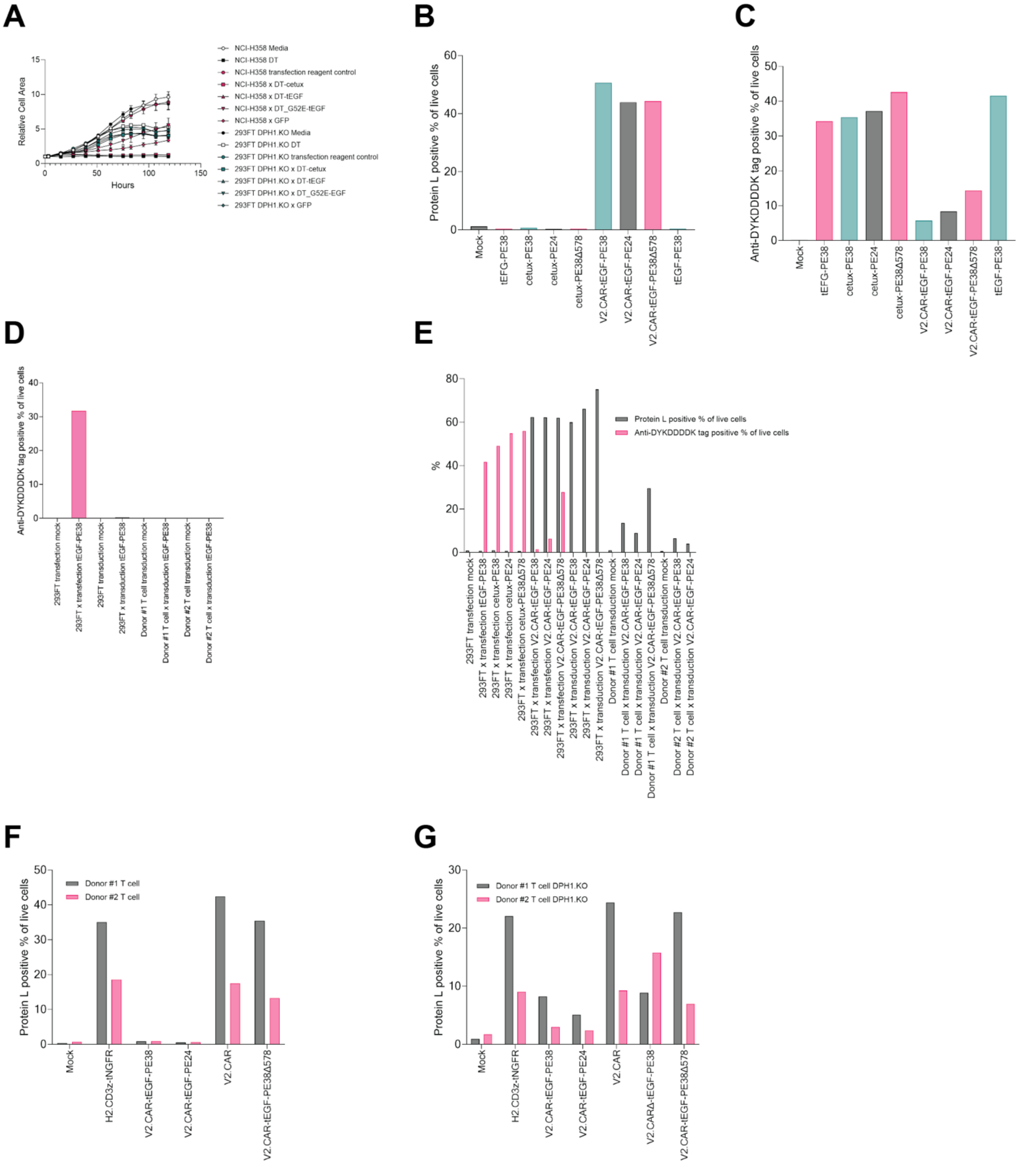


**Supplementary Figure S14: Limitations of functionality of ADPRTs in primary human T cells.** (A) Non-resistant NCI-H358 cells or 293FT cells engineered for ADPRT resistance were transfected with immunotoxin or control constructs. Cell growth was measured over time. (B, C) 293FT cells engineered for ADPRT resistance were transfected with constructs and then assayed by flow cytometry for linked CAR marker expression with cognate tetramer (B) or intracellular FLAG-tag expression (C). (D, E) 293FT DPH1.KO cells or ADPRT resistant primary human T cells were transduced or transfected with immunotoxin constructs. Expression of intracellular FLAG-tag or expression of CAR marker were assayed. (F, G) Unmodified primary human T cells (F) or T cells engineered for ADPRT resistance (G) were transduced with immunotoxin constructs with linked CAR marker. Cells were then assayed for linked CAR marker expression by flow cytometry with protein L staining.

### Supplementary Figure S15

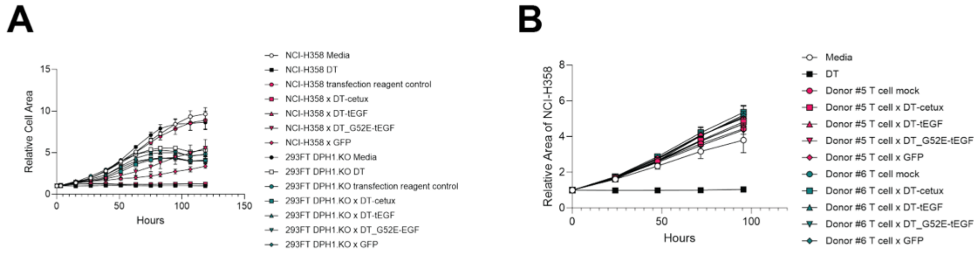


**Supplementary Figure S15: Restricted utility of alternative ADPRT constructs in primary human T cells.** (A) NCI-H358 cells or 293FT DPH1.KO cells were transfected with control constructs or immunotoxin constructs incorporating DT domains. Longitudinal cell growth was measured by microscopy. (B) ADPRT resistant primary human T cells were transfected by electroporation with immunotoxin or control constructs and cultured across a transwell barrier from NCI-H358 target cells. Area of NCI-H358 target cells was measured over time with live cell microscopy.

### Supplementary Figure S16

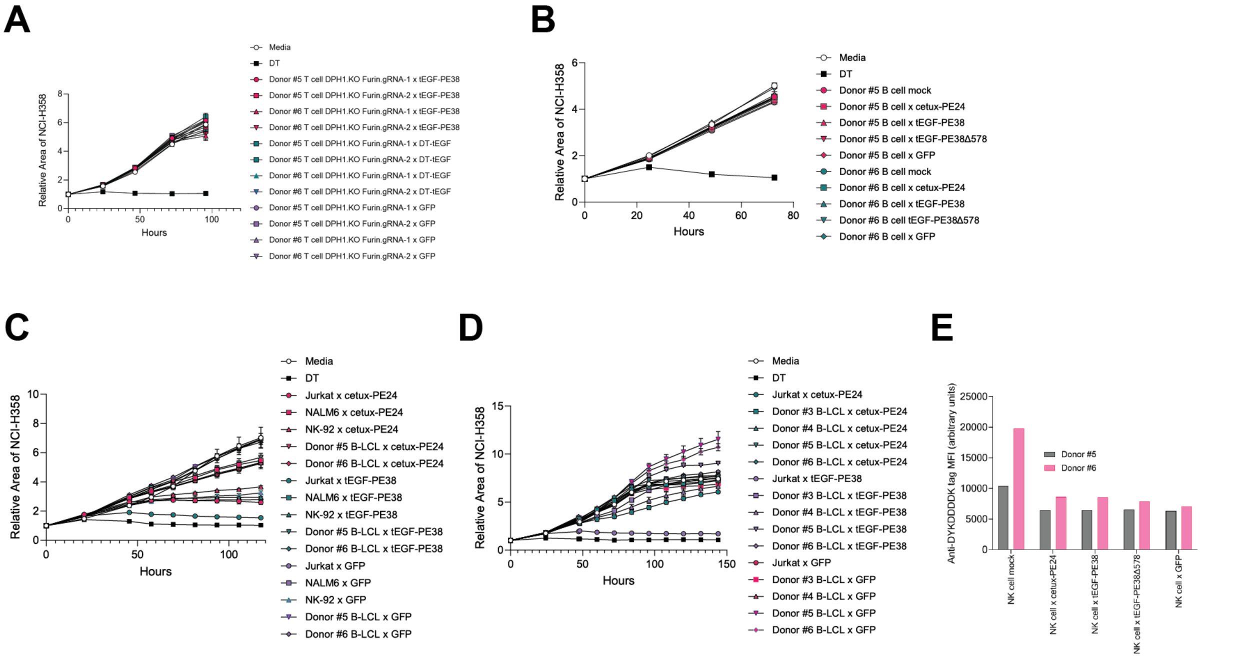


**Supplementary Figure S16: Limited ADPRT expression in alternative primary human cell types.** (A) Primary human T cells were engineered to knock out both *DPH1* and *FURIN* genes. Cells were then transfected by electroporation with immunotoxin constructs and co-cultured in transwell plates with NCI-H358 target cells. NCI-H358 cells were measured longitudinally. (B) Primary human B cells were engineered for ADPRT resistance, transfected with ADPRT constructs, and cultured in transwell plates with NCI-H358 target cells. NCI-H358 cells were quantified across time. (C, D) Jurkat, NALM6, or NK-92 cell lines or human B-LCL were engineered for ADPRT resistance. They were then transfected with immunotoxin constructs and co-cultuered with NCI-H358 cells. Target cell growth was measured over time. (E) Primary human NK cells were engineered for ADPRT resistance and then transfected with immunotoxin constructs. Expression of a linked FLAG-tag was measured by intracellular flow cytometry.
